# A nociceptive amygdala-striatal pathway for chronic pain aversion

**DOI:** 10.1101/2024.02.12.579947

**Authors:** Jessica A. Wojick, Alekh Paranjapye, Juliann K. Chiu, Malaika Mahmood, Corinna Oswell, Blake A. Kimmey, Lisa M. Wooldridge, Nora M. McCall, Alan Han, Lindsay L. Ejoh, Samar Nasser Chehimi, Richard C. Crist, Benjamin C. Reiner, Erica Korb, Gregory Corder

**Affiliations:** Dept. of Psychiatry, Perelman School of Medicine, University of Pennsylvania, Philadelphia, PA, USA; Dept. of Neuroscience, Mahoney Institute for Neurosciences, Perelman School of Medicine, University of Pennsylvania, Philadelphia, PA, USA; Dept. of Biology, School of Arts and Sciences, University of Pennsylvania, Philadelphia, PA, USA; Department of Genetics, Perelman School of Medicine, University of Pennsylvania, Philadelphia, PA, USA; Epigenetics Institute, Perelman School of Medicine, University of Pennsylvania, Philadelphia, PA, USA

## Abstract

The basolateral amygdala (BLA) is essential for assigning positive or negative valence to sensory stimuli. Noxious stimuli that cause pain are encoded by an ensemble of *noci*ceptive BLA projection neurons (BLA^*noci*^ ensemble). However, the role of the BLA^*noci*^ ensemble in mediating behavior changes and the molecular signatures and downstream targets distinguishing this ensemble remain poorly understood. Here, we show that the same BLA^*noci*^ ensemble neurons are required for both acute and chronic neuropathic pain behavior. Using single nucleus RNA-sequencing, we characterized the effect of acute and chronic pain on the BLA and identified enrichment for genes with known functions in axonal and synaptic organization and pain perception. We thus examined the brain-wide targets of the BLA^*noci*^ ensemble and uncovered a previously undescribed *noci*ceptive hotspot of the nucleus accumbens shell (NAcSh) that mirrors the stability and specificity of the BLA^*noci*^ ensemble and is recruited in chronic pain. Notably, BLA^*noci*^ ensemble axons transmit acute and neuropathic *noci*ceptive information to the NAcSh, highlighting this *noci*ceptive amygdala-striatal circuit as a unique pathway for affective-motivational responses across pain states.

## Introduction

It is critical for survival to quickly and appropriately adapt to environmental stimuli or internal states that signal threats of bodily harm—the primary function of the nociceptive nervous system that produces the perception of pain (1). Aversion to pain, or the unpleasantness of the experience, relies on nociception to engage negative valence neural circuits in the brain that work in concert to initiate protective escape and avoidance behaviors (2). Chronic pain is a disease state wherein intensely unpleasant and persistent pain is perceived, despite any recuperative behavioral actions, suggesting the maladaptive engagement of valence neural circuitries.

The basolateral subnucleus of the amygdala (BLA) has been investigated for its valence-assigning role over decades (3), with debate about the specific cell types and anatomical architectures that transform sensory information into positive or negative valence signals to be distributed across the brain (4–11). We recently demonstrated that the BLA contains a functional subpopulation of negative valence neurons encoding *noci*ception that is essential for the emotionally aversive aspect of *noci*ception, regardless of sensory modalities and distinct from appetitive processes—the BLA^*noci*^ ensemble (12). Inhibition or activation of the BLA^*noci*^ ensemble decreased or augmented *noci*fensive behavior, respectively, specifically pertaining to aversive affective-motivational responses, such as attending to injured tissue, escape, and future avoidance of noxious events (12, 13). Importantly, using a cross-day tracking analysis of the calcium (Ca^2+^) activity of individual BLA neurons, we previously observed that same individual BLA neurons responsive to acute noxious stimuli respond to formerly innocuous stimuli following neuropathic injuries (12). Thus, the BLA^*noci*^ ensemble represents a unique neural circuit for understanding how adaptive behaviors emerge from precise valence neural ensembles, but also for translational analgesic efforts that might reduce chronic pain aversion by targeting this stable *noci*ceptive BLA ensemble.

To date, the functional stability of the BLA^*noci*^ ensemble to facilitate aversive behaviors during acute and chronic pain has not been directly explored. Additionally, we do not know the brain-wide projection map or genetic composition of this functional ensemble, nor how acute and chronic pain alters these characteristics. Single cell transcriptomic profiling of BLA cell-types recently uncovered unique neuronal subpopulations that express previously described, single gene markers of putative valence populations (6, 14– 19). Others have dissected BLA valence populations based on projection target (7, 10), but it is unknown if *noci*ceptive information is transmitted to these downstream brain regions. BLA principal neurons send long-range glutamatergic projections to many affective brain regions, including the nucleus accumbens (NAc) (20, 21), a striatal structure important for driving aversive and appetitive behavior (22). In addition, recent work demonstrated that the BLA transmits negative valence information to the NAc (10, 23–25), despite previous evidence suggesting it is a purely appetitive circuit (7, 21, 26, 27). Further, dysfunction of the NAc has been implicated in chronic pain disorders in preclinical and clinical studies (28–31), particularly in the limbic NAc shell (NAcSh) (32–34). While the role of the BLA projections to the NAcSh in appetitive motivation have been well-characterized (35–39), little is known about the role of this circuit in processing unconditioned *noci*ceptive information.

The NAcSh has canonically been viewed as primarily important for reward and appetitive motivation. However, recent studies have shown that the anatomical position (40, 41), genetic identity (39, 42–44), and endogenous opioid expression (45, 46) of NAcSh neurons may underlie distinct and opposing behavioral functions. Clinical positron emission tomography imaging studies and preclinical electrophysiological and behavioral studies have highlighted the importance of NAcSh neurons in chronic pain (22, 29, 32, 33, 42). Despite this, the location of neurons processing *noci*ception in the NAcSh has not been identified nor has a direct link between functional *noci*ceptive circuits in the BLA and NAcSh been explored.

In this study, we sought to understand how the presence of pain, both acutely and chronically, impacts the BLA in terms of behavioral function, transcriptional identity, and downstream projection activity. We found that the BLA^*noci*^ ensemble is a functionally unique subpopulation of negative valence BLA neurons that is required for the expression of acute and chronic pain-related aversive behaviors. We used single nucleus RNA-sequencing to define the transcriptomic changes that accompany acute and chronic pain in the BLA and identified multiple enriched genes implicated in pain and negative affect and genes associated with synaptic transmission. Thus, we examined the downstream *noci*ceptive brain regions of the BLA^*noci*^ ensemble and identified a previously undescribed *noci*ceptive hotspot of the NAc in the posterior medial dorsal NAcSh. This *noci*ceptive NAcSh hotspot mirrors the *noci*ceptive stability and valence specificity of the BLA^*noci*^ ensemble, suggesting it may be an important node in the affective *noci*ceptive brain network. Therefore, we tested whether there is *noci*ceptive information transmission from the BLA to the NAcSh using axon Ca^2+^ activity recording. In agreement with our transcriptomic dataset suggesting heterogeneity in functional activity within genetic populations, we found that, despite previous characterizations of *Rspo*2+ BLA neurons as the purported negative valence population, *Rspo*2+ BLA axons transmit mixed valence information to the NAcSh. In contrast, BLA^*noci*^ ensemble axons display *noci*ceptive-specific information transmission to the NAcSh, including signals related to neuropathic pain allodynia. Thus, multilevel examination of a BLA^*noci*^ ensemble projecting to the NAcSh allowed us to refine the understanding of negative valence processing between the amygdala and ventral striatum. Furthermore, this work highlights this amygdala-striatal circuit as a potential target for novel treatments to lessen chronic pain aversion.

## Results

### Acute and neuropathic nociception engages a shared valence ensemble in the BLA

We previously found a *noci*ceptive ensemble of BLA neurons (BLA^*noci*^ ensemble) that encodes acute and chronic *noci*ceptive stimuli regardless of sensory modality but does not encode non-*noci*ceptive aversive nor appetitive stimuli (12). However, it remains unknown how the stability of the BLA^*noci*^ ensemble across pain states translates to function. Thus, our goal was to determine the cellular stability and valence selectivity of *noci*ception within individual BLA neurons, across time, valence assignment, and different pain models. To do this, we leveraged the updated TRAP2 mouse (*Fos*-FOS-2A-iCre^ERT2^) crossed with the Ai9 tdTomato fluorescent reporter mouse line (TRAP2:tdTomato) to capture functional ensembles across the brain (47). We genetically captured functionally *noci*ceptive neurons using a *noci*TRAP protocol: application of noxious 55°C water droplets to the left hindpaw (10 drops [∼50 µl/drop], once per 30 sec, over 10 min) and administered 4-hydroxytamoxifen (4-OHT; 40 mg/kg, s.c.). This *noci*TRAP protocol initiates genetic recombination and permanent expression of tdTomato in *noci*ceptive cells throughout the nervous system (**Fig. 1A, B**). Two weeks later, we exposed the *noci*TRAP2:tdTomato mice to a second noxious 55°C water stimulus to drive expression of the IEG FOS (*noci*FOS) and examined the colocalization of *noci*FOS with *noci*TRAP neurons (**Fig. 1A, B**). We found that *noci*TRAP and *noci*FOS neurons spread across the anterior-posterior axis of the lateral and basal subnuclei of the BLA, with 43.92% co-localization, suggesting stability of the BLA^*noci*^ ensemble to repeatedly encode *noci*ception (**Fig. 1C-E**). This degree of neural re-activation between two temporally distant, similar stimuli aligns with previous colocalization reports using the TRAP2 mouse (48–51). Furthermore, we did not find that basal, non-sensory evoked FOS levels (*home-cage*FOS) colocalize to the same degree as a second *noci*ceptive stimulus (**Fig. 1F**). Importantly, we next tested the valence specificity of *noci*TRAP by comparing the colocalization of neurons captured during an appetitive mating opportunity (*mate*TRAP), wherein male TRAP2:tdTomato mice mounted a novel female mouse and intromissions occurred. Compared to *noci*TRAP, we observed little colocalization between *mate*TRAP neurons and *noci*FOS neurons, confirming previous reports that valence processes are partially segregated between distinct neuronal ensembles (4, 6–8, 10, 12, 27) (**Fig. 1G**).

**Figure 1.**
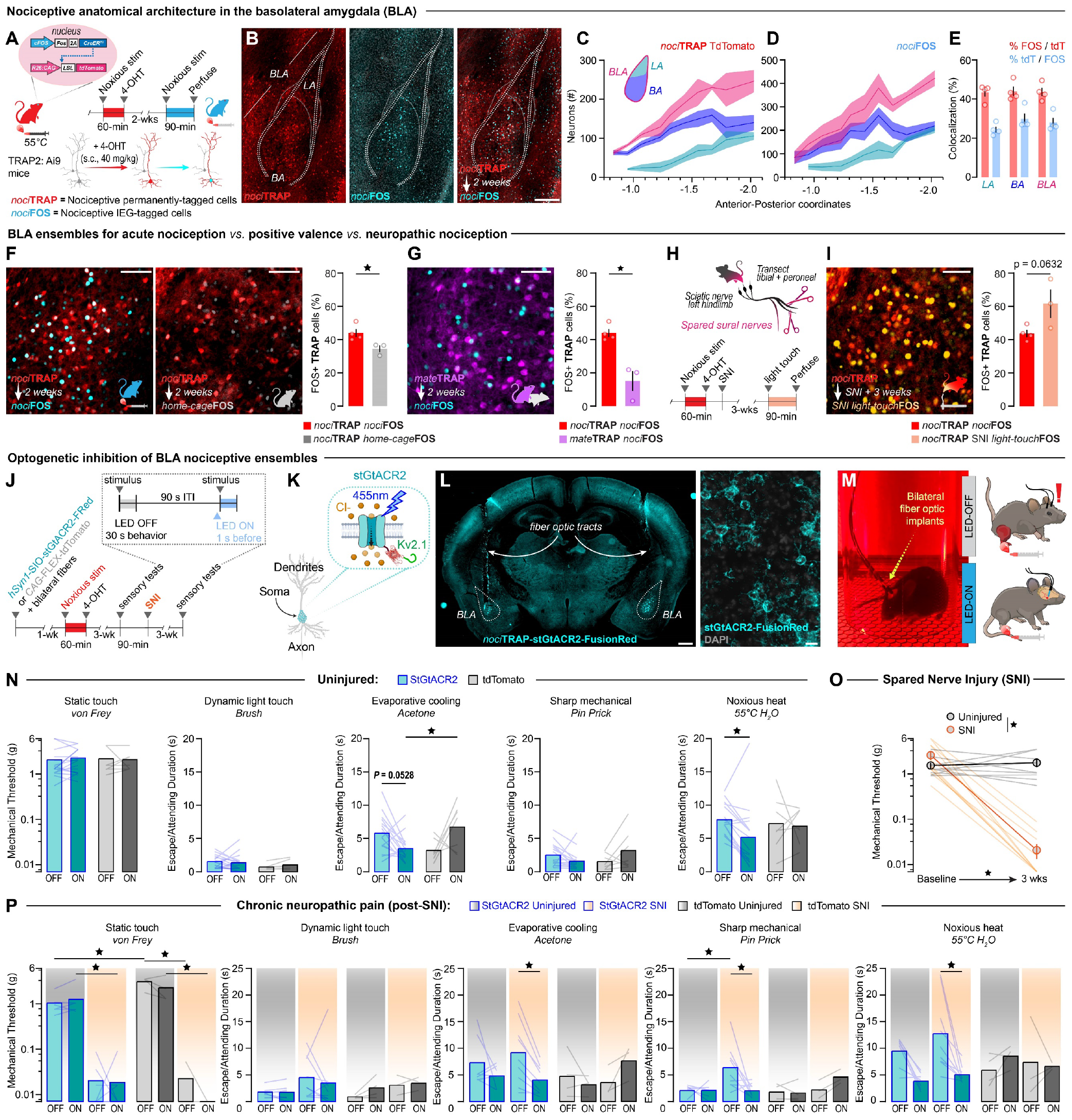
Inhibition of the BLA^*noci*^ ensemble reduces pain aversion in acute and chronic pain. **(A)** *Noci*ceptive TRAP (*noci*TRAP) protocol. **(B)** Representative images of neurons captured by *noci*TRAP tdTomato (red) and *noci*FOS (blue) with quantification of **(C)** *noci*TRAP **(D)** and *noci*FOS across the anterior-posterior axis of the basolateral amygdala (BLA), lateral amygdala (LA), and basal amygdala (BA). Scale: 250 μm. **(E)** Quantification of the colocalization of *noci*TRAP and *noci*FOS in neurons across the LA, BA, and BLA (n = 4). **(F)** Representative images of *noci*TRAP colocalization with *noci*FOS (blue) vs *home-cage*FOS (grey) and quantification (Two-tailed unpaired *t* test, p =0.0379, n = 4 *noci*FOS, n = 3 *home-cage*FOS). Scale: 100 μm. **(G)** Colocalization of *mate*TRAP (purple) and *noci*FOS (blue). Scale: 100 μm. Quantification of colocalization of *noci*FOS with neurons captured by *mate*TRAP relative to *noci*TRAP. (Two-tailed unpaired *t* test, p =0.0042, n = 4 *noci*TRAP, n = 3 *mate*TRAP). **(H)** Schematic and timeline of spared nerve injury (SNI) model of chronic neuropathic pain. **(I)** Colocalization of uninjured *noci*TRAP (red) and light-touch after SNI FOS (orange). Scale: 100 μm. Quantification of colocalization of *light-touch*FOS in animals than underwent SNI relative to a second acute *noci*ceptive stimulus in uninjured mice. (Two-tailed unpaired *t* test, p =0.0632, n = 4 *noci*FOS, n = 3 SNI *light-touch*FOS). **(J)** Timeline of optogenetic inhibition experiment. **(K)** Schematic of inhibitory opsin. **(L)** Histological confirmation of bilateral expression of stGtACR2 in BLA^*noci*^ ensemble with bilateral fiber optics above. Scale: 500 μm (left image), 250 μm (right image). (**M)** Image of behavioral testing setup. **(N)** Optogenetic inhibition of the BLA^*noci*^ ensemble decreases behavioral responding to noxious stimuli compared to a pre-stimulus baseline. (One-way repeated-measure ANOVA with Bonferroni; acetone: main effect of interaction: p = 0.0032, light ON stGtACR2 vs tdTomato: p = 0.0134; 55°C water: main effect of mouse: p = 0.0161, stGtACR2 light OFF vs light ON: p = 0.0249; n = 16 StGtACR2 (8 male), n = 7 tdTomato (2 male)). **(O)** Mice that received a SNI to induce chronic neuropathic pain showed hypersensitivity three weeks post-SNI. (Two-Way repeated-measure ANOVA with Bonferroni, main effect of interaction: p = 0.0007, SNI group baseline vs. 3-weeks post-SNI: p <0.0001, 3-weeks post-SNI uninjured vs. SNI: p = 0.0029). **(P)** Increased responding to innocuous and noxious stimuli was reduced during inhibition of the BLA^*noci*^ ensemble compared to pre-stimulus baselines. (Three-way ANOVA with Tukey; von Frey: main effects of virus (p = 0.0099), injury (p <0.0001), interaction of virus and injury (p = 0.0095), LED OFF Uninjured StGtACR2 vs tdTomato: p = 0.0003, LED ON StGtACR2 uninjured vs SNI: p = 0.0101, LED OFF tdTomato uninjured vs SNI: p <0.0001, LED ON tdTomato uninjured vs SNI: p = 0.0012; acetone: main effect of interaction: p = 0.0483, SNI StGtACR2 LED ON vs LED OFF: p = 0.0346; 55°C water: main effects of LED (p =0.0341) and interaction of LED and virus (p = 0.0067), StGtACR2 SNI LED OFF vs LED ON: p = 0.0089). (Pin prick: Mixed effects analysis with Tukey, main effect of interaction: p = 0.0261, StGtACR2 LED OFF uninjured vs SNI: p = 0.0143, StGtACR2 SNI LED OFF vs LED ON: p = 0.0060). N = 7 StGtACR2 uninjured (4 male), n = 9 StGtACR2 SNI (4 male), n = 4 tdTomato uninjured (2 male), n = 3 tdTomato SNI (all female)). BLA = basolateral amygdala.

Previously, single-cell resolution Ca^2+^ imaging of the BLA revealed that acute and chronic pain are encoded by the same BLA neurons. Here, we confirmed this finding by examining the overlap of *noci*TRAP and FOS activity in a chronic neuropathic pain state. To induce chronic pain, we performed the spared nerve injury (SNI) model of chronic neuropathic pain on *noci*TRAP:tdTomato mice (52) (**Fig. 1H**). Three weeks after SNI, we exposed the injured left hindpaw to a static light-touch with a 0.16 gram von Frey filament to evoke FOS (*light*-*touch*FOS) as a model of mechanical allodynia. We found that light-touch in an SNI state reactivated a similar percentage of *noci*TRAP neurons compared to *noci*TRAP mice receiving a second noxious stimulus in an uninjured state. This suggests that the BLA^*noci*^ ensemble is stably responsive across the acute to chronic pain transition (**Fig. 1I**).

### Acute and chronic pain aversion requires a common BLA nociceptive neural ensemble

Our current and previous observations of a stable and valence-specific BLA^*noci*^ ensemble for acute and chronic pain-related processes raises the question of whether the same neurons active in acute pain are required for chronic pain. We tested this critical question using an optogenetic approach, expressing the soma-targeted blue-light inhibitory opsin, stGtACR2-FusionRed (53), or a control tdTomato fluorophore, bilaterally in BLA^*noci*^ neurons of *noci*TRAP2 mice. We examined stimuli-evoked reflexive and affective-motivational (i.e., nocifensive) behaviors before and during inhibition of BLA^*noci*^ neurons (**Fig. 1J-M**). We demonstrated that, in support of our previous chemogenetic results (12), BLA^*noci*^ ensemble inhibition did not affect sensory thresholds, but decreased *noci*fensive behavior to noxious stimuli in uninjured mice (**Fig. 1N**). Furthermore, in a real-time place preference assay, inhibition of the BLA^*noci*^ ensemble was not reinforcing (**Fig. S1A**). Interestingly, chemogenetic activation of the BLA^*noci*^ ensemble in the absence of injury caused a conditioned place aversion in female mice, but had no effect on *noci*ception-related behavioral responses (**Fig. S2**). This suggests that activity of the BLA^*noci*^ ensemble can drive negative affective behavior but does not further potentiate *noci*ceptive behavior when this ensemble is already active.

Our next goal was to determine whether these same neurons within the BLA^*noci*^ ensemble also causally contribute to chronic neuropathic pain behavior. We performed the same optogenetic inhibition protocols in the same mice 3 weeks after either SNI or no-injury. SNI mice demonstrated neuropathic hypersensitivity compared to their pre-injury responses and to uninjured controls (Fig. 1O). While inhibition did not alter mechanical allodynia thresholds, we observed decreased allodynia and hyperalgesia behavior to acetone, pin prick and 55°C water droplets during blue-light inhibition in SNI *noci*TRAP:stGtACR2 mice but not SNI *noci*TRAP:tdTomato controls. (**Fig. 1P**). This demonstrates that chronic neuropathic pain relies on the same BLA^*noci*^ ensemble circuitry as acute pain and engagement of this ensemble is stable across the acute to chronic pain transition. No effect was observed on place preference for the SNI *noci*TRAP:stGtACR2 mice or other groups, suggesting that inhibition of the BLA^*noci*^ ensemble is not engaging motivational or reward-related neural circuitry (**Fig. S1B**). Together, these results demonstrate that the BLA^*noci*^ ensemble is a functionally-stable and valence-specific ensemble of neurons facilitating *noci*fensive behaviors.

### Transcriptomic atlas of the nociceptive amygdala

Our histological and optogenetic data, in addition to our prior single-cell Ca^2+^ imaging of the BLA (12), confirms that the BLA^*noci*^ ensemble is a functionally distinct set of neurons capable of dictating specific behavior responses. However, the molecular signatures that distinguish this *noci*ceptive ensemble remain undefined. The genetic identity of the BLA^*noci*^ ensemble has not been previously examined beyond a few genes, such as *Slc171a7* (VGLUT1), identifying the ensemble as glutamatergic, and *Rspo2*, a marker for the putative negative valence BLA ensemble (6, 8, 12). Therefore, to define the transcriptomic signatures of acute and chronic pain in an unbiased manner, we used single nucleus RNA-sequencing (snRNAseq) to transcriptionally profile the BLA. We collected amygdalar tissue (right hemisphere only) from uninjured or SNI mice immediately following (∼5 min) either no stimulus, a light-touch, or a noxious 55°C water droplet stimulation of the left hindpaw (**Fig. 2A**). All tissue samples were collected and fully processed on the same day to avoid batch effects. Analysis of nuclei from uninjured groups identified 30 cell-type clusters, including 18 neuronal clusters, from 72,125 nuclei (**Fig. 2B**). Nuclei from all stimulus treatments were equally distributed across the 30 clusters (**Fig. 2C, D**). Based on known cell-class-defining marker genes and differentially expressed genes (DEGs), we parsed the clusters into nine GABAergic neuron clusters, six glutamatergic neuron clusters, three neuron clusters that labeled putatively non-amygdalar brain regions (*e*.*g*., caudate putamen), and twelve non-neuronal clusters (*e*.*g*., microglia, astrocytes, etc.) (**Fig. 2D-F**). With our large-scale amygdalar transcriptomic atlas, we next homed in on two subnuclei that have been strongly implicated in pain—the central amygdala (CeA) and the BLA (54, 55). From the classified neuronal pool, based on published markers and the Allen Brain in situ hybridization database (**Fig. S3**), we initially defined BLA subclusters as nuclei in the ≥75th percentile for *Slc17a7*, and putative CeA as the ≥75th percentile for Gad1 and/or *Gad2* (and *Slc17a6*- and *Slc17a7*-). Further clustering yielded 13 inhibitory subclusters, eight putative BLA subclusters, one intercalated cell (ITC9) cluster, and one endopiriform cortex (Endopiro5) cluster (**Fig. 2G, H, Fig. S3**). Subsequently, we compared several published marker genes for unique cell types involved in valence processes and *noci*ception within the BLA and CeA; namely, *Rspo2* and *Fezf2* for BLA valence ensembles (6, 8, 10, 12), and Prkcd (PKCδ), Sst (Somatostatin), and Calcrl (CGRP receptor) for CeA valence/*noci*ception ensembles (56–63) (**Fig. S3**). Transcriptional profiles did not adequately distinguish the BLA GABAergic cell-types (e.g., somatostatin, parvalbumin, vasoactive intestinal peptide interneurons) from the CeA GABAergic cell-types expressing the same marker genes; thus, we have labeled all *Gad1*+ and *Gad*2+ nuclei as “inhibitory cell” clusters rather than CeA, and no further analyses are conducted on these cells within this study.

**Figure 2.**
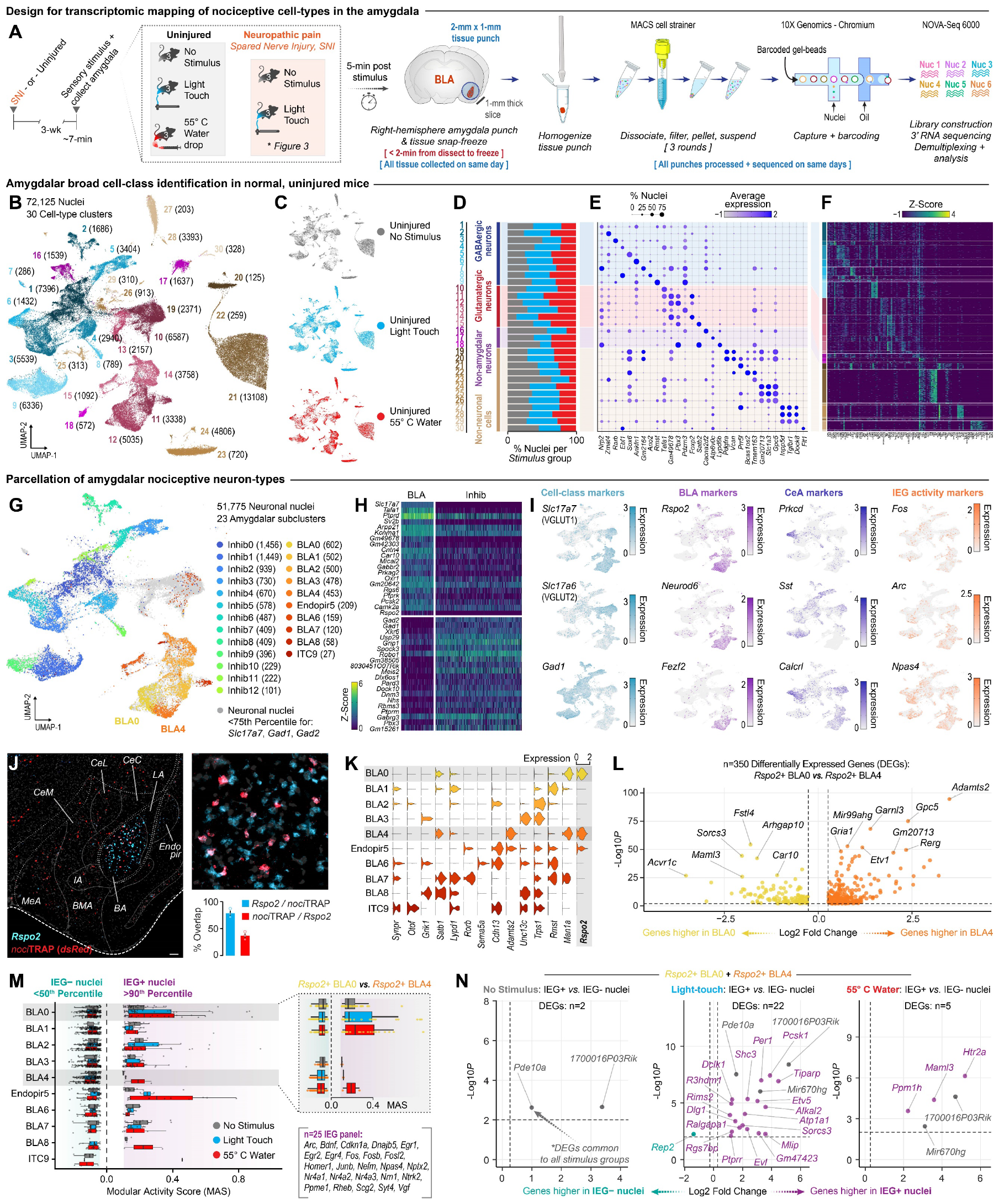
The BLA^*noci*^ ensemble is a functional subpopulation of *Rspo2* BLA neurons. (A) Timeline, experimental design, and methods. (B) UMAP of all nuclei (n = 72,125) captured by punches around the amygdala in 30 unique clusters. (C, D) Contribution of nuclei from various stimulation conditions to the total UMAP and cell types. (E, F) Dotplot and heatmap displaying a top gene differentiating the 30 cell clusters. (G) UMAP of amygdalar neuron nuclei (n = 51,775) showing 13 inhibitory (*Gad1*+ *or Gad2*+) and 10 BLA (*Vglut1*+) putative clusters. (H) Heatmap displaying top genes differentiating the 23 neural clusters. (I) Feature plots of select genes for cell classes, published gene markers for valence/*noci*ception in the BLA and CeA, and immediate early genes (IEGs). (J) Representative 4X and 20X fluorescent images and quantification of RNAscope fluorescent *in situ* hybridization (FISH) showing that BLA^*noci*^ neurons largely express *Rspo2* but are a subpopulation of all BLA *Rspo2*+ neurons. Scale: 100 µm. (K) Violin plot displaying expression of genes identified in past BLA RNA sequencing, including expression of *Rspo2*, by subcluster. (L) Volcano plot of 350 differentially expressed genes (DEGs) between *Rspo2*+ BLA clusters 0 and 4. Yellow dots indicate genes enriched in the BLA subcluster and orange dots indicate genes enriched in the BLA4 subcluster. (M) IEG modular activity scores of 10 putative BLA clusters across stimulation conditions segregated by percentile thresholds. (N) Volcano plots of BLA clusters 0 and 4 displaying DEGs upper stimulation condition enriched in the IEG+ or IEG-nuclei. Purple dots = DEGs only identified within that condition. Grey dots = shared DEGs across conditions.

Next, we mapped several IEGs as proxies for stimulus-evoked neuronal activity and observed expression across the UMAP space including within the *Rspo*2+/Slac17a+ regional clusters (**Fig. 2I**). As *Rspo*2+ BLA neurons have been implicated in negative valence processing (6, 8), we explored the colocalization of *Rspo2* in *noci*TRAP BLA neurons and found that 78.63% of *noci*TRAP BLA neurons express *Rspo2* (**Fig. 2J**). Therefore, we examined the expression of *Rspo2* and other genes among the putative-BLA subclusters and identified three subclusters that express *Rspo2*: BLA0, BLA4, and a third cluster with low *Rspo2* (defined as Endpiro5) (**Fig. 2K**). Upon further investigation of the Allen Brain in situ hybridization database based on DEG analysis of these three clusters, we confirmed that genes enriched in Endopir5 largely map to the endopiriform cortex, while additional BLA0 and BLA4 enriched genes specifically localize to the BLA (**Fig. 2K, Fig. S3D, E**). Related to prior genetic identities for valence ensembles, we do not observe co-expression of *Fezf2* nor *Ppp1r1b* within our *Rspo*2+ BLA0 and BLA4 subclusters (**Fig. S3A**). We identified 350 DEGs between the *Rspo*2+ BLA0 and BLA4 subclusters (**Fig. 2L**), including many genes implicated in negative affective and pain behavior, such as *Sorcs3, Fstl4, Garnl3* (64–66). Together, these data define two *Rspo*2+ BLA subclusters with distinct gene expression profiles from each other, both of which include genes linked to pain and negative affective behavior.

Next, we sought to identify BLA^*noci*^ neurons using transcriptional profiles of *Rspo*2+ BLA cells. Notably, the BLA^*noci*^ neurons are a subpopulation of the total *Rspo*2+ BLA neurons, comprising 36.86% (**Fig. 2J**). To identify the *noci*ceptive BLA cell-types, we examined the modular expression of a panel of 25 IEGs (67) against randomly selected background expression between our three stimuli conditions (no stimulus, light touch, and noxious 55°C water droplets) across BLA subclusters. Interestingly, the BLA subclusters displayed variable IEG activity in response to the stimuli, including to the noxious 55°C water droplets. This suggests that despite genetic similarities that grouped nuclei into the subclusters, there is heterogenous functional activity of BLA nuclei, even within the *Rspo*2+ BLA subclusters 0 and 4 (**Fig. 2M**).

Individual analyses of BLA0 and BLA4 did not provide sufficient cell numbers to confidently identify DEGs between IEG+ and IEG-cells. However, across the two *Rspo*2+ BLA subclusters combined, we identified the genes that are differentially expressed across stimulation conditions (**Fig. 2N**, with purple indicating genes only identified as differentially expressed within that condition and grey indicating shared DEGs across conditions). Three of these genes (*Ppm1h, Maml3*, and *5Htr2a*) are uniquely upregulated in response to noxious stimulation. Interestingly, other genes (*Pde10a, Alkal2*) that are found in multiple IEG+ groups or in response to non-noxious stimuli are implicated in pain and negative affect (68, 69), suggesting that the highly active BLA0/4 neurons express greater levels of genes related to pain processing (Fig. 2N). These findings define a BLA cell population that is highly responsive to nociceptive stimuli, providing the first transcriptomic classification of pain-active neurons in the amygdala.

### Chronic neuropathic pain leaves a transcriptomic imprint on nociceptive BLA cell-types

While our behavioral and anatomical data suggests that the BLA is uniquely important for the expression of acute and chronic pain, the genetic changes that accompany pain chronification in the BLA remain unclear. To test this, we used snRNAseq as above to compare the BLA from uninjured and three weeks post-SNI mice (**Fig. 2A**). From 73,512 neuronal nuclei across four injury and stimulus conditions, we identified the same 10 glutamatergic cell-types (**Fig. 3A**), which expressed a similar pattern of marker genes defining the amygdalar cell-types from only the uninjured only mouse cohorts in Figure 2G (**Fig. 3B, C**).

**Figure 3.**
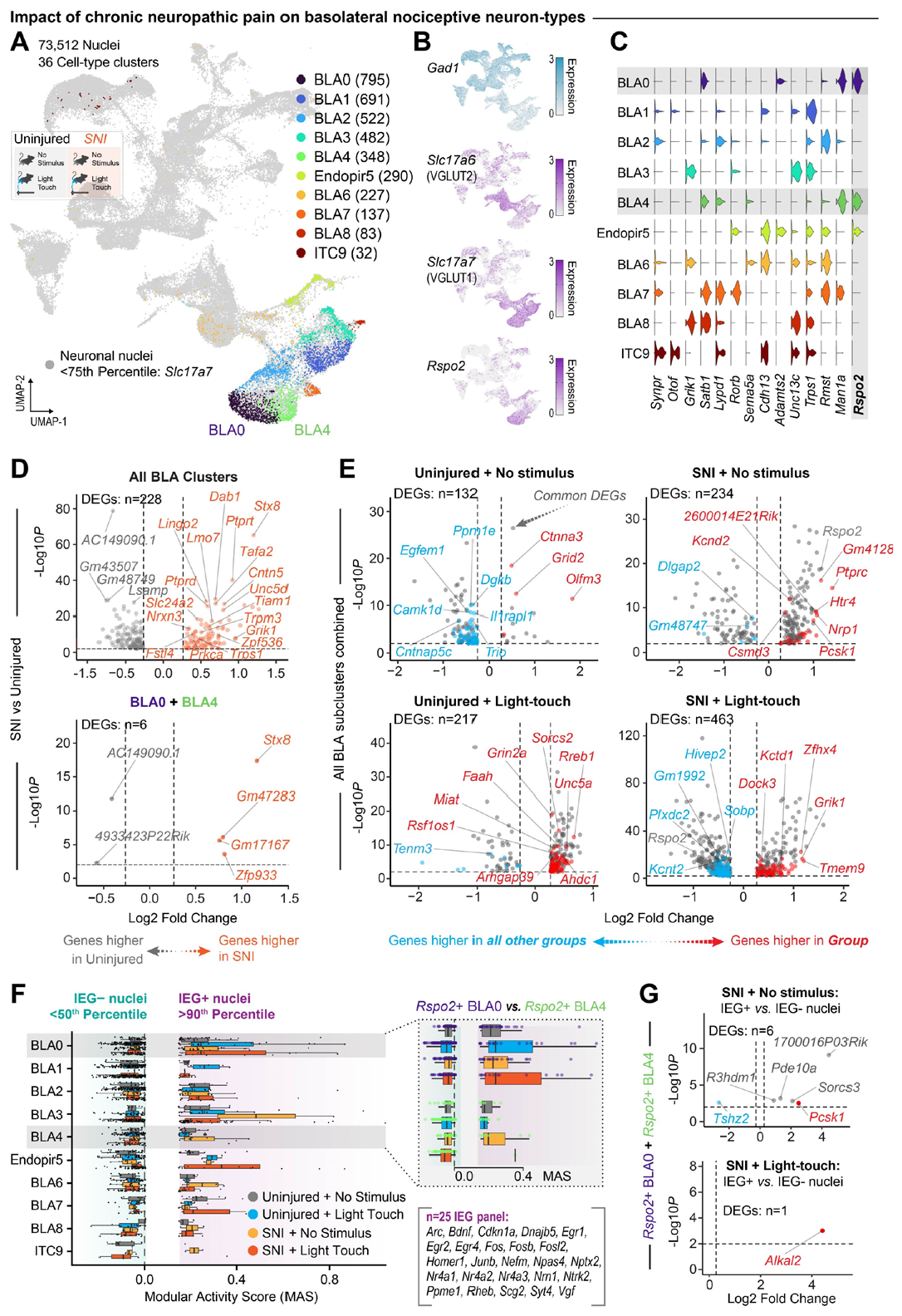
Chronic pain imparts a unique transcriptomic signature on the BLA. (A) UMAP of all neuronal nuclei (n = 73,512) across uninjured and chronic neuropathic pain (SNI) conditions identifying 36 unique cell-type clusters. (B) Feature plots of genes for cell classes and for *Rspo2*. (C) Violin plot displaying expression of genes identified in past BLA RNA sequencing, including expression of *Rspo2*, by subcluster. (D) Volcano plots of all BLA subclusters combined (top) or subclusters BLA0 and BLA4 combined (bottom) displaying DEGs in all nuclei from SNI mice compared to nuclei from uninjured mice. Orange dots indicate genes enriched in SNI conditions, gray dots indicate genes enriched in uninjured conditions. (E) Volcano plots of all BLA subclusters combined displaying DEGs unique across injury and stimulation conditions. Red dots indicate genes enriched in that condition, blue dots indicate genes enriched in all other conditions, and grey dots indicate shared DEGs across conditions. (F) IEG modular activity scores of 10 putative BLA clusters across stimulation conditions. (G) Volcano plots of *Rspo2*+ BLA subclusters 0 and 4 displaying DEGs unique between IEG+ and IEG-nuclei in SNI. Red dots indicate genes enriched in that condition, blue dots indicate genes enriched in all other conditions, and grey dots indicate shared DEGs across conditions.

We identified the genes that are differentially expressed in a chronic neuropathic pain state across the eight putative BLA subclusters and identified 228 DEGS including multiple upregulated genes in the SNI condition that have been implicated in *noci*ception (*Stx8, Nrxn3, Slc24a2, Prkca, Ptprd, Unc5d, Lmo7, Tiam1*, and *Trpm3*) or other amygdalar processes involving fear and negative affect (*Fstl4, Zpf536, Trps1, and Dab1*, and *Grik1*) (65, 70–82) (**Fig. 3D**). We examined the function of these DEGs using gene ontology analysis and found that the majority of the DEGs relate to axon generation and guidance, and synaptic transmission (**Fig. S4**). This suggests that the major impact of chronic pain on the BLA is on the functional connectivity of the BLA to downstream brain regions. To specifically examine the impact of chronic pain on the negative valence BLA neurons, we focused on clusters BLA0 and BLA4 due to their high expression of *Rspo2* (**Fig. 3C**). Within these two clusters, we found that SNI resulted in differential expression of only six genes, including Stx8 and Gm47283, which have been implicated in the expression of chronic pain behaviors (70, 83, 84) (**Fig 3D**). Next, we compared the DEGs unique to each of the four injury and stimulation conditions across all BLA neurons, and identified many genes that are differentially expressed following SNI (**Fig. 3E)**. Genes uniquely upregulated or downregulated in these conditions, relative to no-injury control nuclei, are colored red or blue, respectively; while the neuropathic-altered DEGs common to both SNI + No stimulus and SNI and light-touch are colored gray. Interestingly, while *Rspo2* is upregulated in the SNI + no stimulus condition, it is downregulated in the SNI + light-touch condition (**Fig. 3E**). This indicates that there is differential expression of *Rspo2* in baseline vs. evoked neuropathic pain conditions. To home in on the *noci*ceptive neurons of the BLA, we examined the expression of 25 IEGs (67) in response to no stimulus or a light touch in uninjured or SNI mice to identify the subset of neurons that are active in chronic pain conditions. Similar to an uninjured state, BLA0 and BLA4 neurons displayed variable IEG activity in response to a light touch in an SNI state, suggesting heterogenous changes in baseline activity following SNI and that subsets of each cluster are active under basal conditions and following light-touch in SNI mice (**Fig. 3F**). While individual analysis of BLA0 and BLA4 again did not provide sufficient cell numbers to identify DEGs in IEG+ versus IEG-nuclei, the combined *Rspo*2+ BLA0 and BLA4 subclusters across stimuli in SNI mice identified six DEGs in the SNI + no stimulus condition and one DEG, *Alkal2*, in the SNI light-touch condition (**Fig. 3G**). *Alkal2*, a ligand for the tyrosine kinase receptor, has been previously demonstrated to regulate the behavioral expression of chronic pain (69). This suggests that *Alkal2*-related processes may partially mediate the *noci*ceptive negative valence of chronic pain within the *Rspo*2+ BLA neurons. Together, this transcriptional characterization of the BLA in acute and chronic pain identifies multiple candidate genes that are altered by chronic pain and an enrichment of genes involved in axon regulation and synaptic transmission (**Fig. S4**). This highlights the necessity of examining the effect of chronic pain on the axonal projections of BLA^*noci*^ neurons at distinct downstream brain regions.

### The BLA^noci^ ensemble projects to a nociceptive hotspot in the nucleus accumbens

Single nucleus RNA-sequencing of the BLA suggests that chronic pain changes differential expression of genes related to downstream axon regulation and signaling. Therefore, we next sought to examine the brain-wide targets of the BLA^*noci*^ ensemble to investigate the routes of *noci*ceptive information flow. To begin, we traced axon outputs of the BLA^*noci*^ ensemble using anterograde fluorescent tracing. We transfected the BLA of TRAP2:tdTomato mice with a Cre-dependent viral GFP and captured *noci*ceptive neurons using our *noci*TRAP protocol, which labels all *noci*ceptive neurons throughout the brain with tdTomato and fills the BLA^*noci*^ ensemble with GFP for axon mapping within downstream regions (**Fig. 1A and Fig. 4A**). The viral injection was primarily localized to the BLA, with some minor spread to the adjoining piriform cortex. We then visualized and quantified BLA^*noci*^ ensemble axon density throughout the brain. We observed BLA^*noci*^ ensemble axons in brain regions with known roles in affective processing (*e*.*g*. anterior cingulate cortex, insular cortex, olfactory tubercle, nucleus accumbens, bed nucleus of the stria terminalis, and central amygdala) that contained high numbers of *noci*TRAP:tdTomato neurons, which we used to guide further detailed quantification (**Fig. 4B, C**). Interestingly, relative to the other downstream brain regions, we observed a *noci*ceptive hotspot in the nucleus accumbens shell (NAcSh) (**Fig. 4D, E**). We then mapped the specific location of these *noci*ceptive neurons throughout the NAcSh and found a dense cluster in the medial-dorsal aspect that increased in the posterior coordinates of the NAcSh (**Fig. 4F-H**). The NAcSh *noci*ceptive neurons (NAcSh*noci*) were located in close proximity to BLA^*noci*^ ensemble axons (**Fig. 4G, H**). To further examine the cellular stability and valence selectivity of *noci*ception within individual NAcSh^*noci*^ neurons across different pain models, time, and valence assignment, we performed similar experiments using the TRAP2:tdTomato mouse as in the BLA (**Fig. 1**). We found that *noci*TRAP and *noci*FOS neurons of the NAcSh displayed 44.79% co-localization, suggesting stability of the NAcSh^*noci*^ ensemble to repeatedly encode *noci*ception. Furthermore, we did not find that basal, non-sensory evoked FOS levels in the NAcSh^*noci*^ (*home-cage*FOS) colocalize to the same degree as a second *noci*ceptive stimulus (**Fig. 4I**). Importantly, we next tested the valence specificity of the NAcSh^*noci*^ ensemble by comparing the colocalization of neurons captured during *mate*TRAP. We observed little colocalization between mateTRAP neurons and *noci*FOS neurons (**Fig. 4J**). As the NAcSh has been implicated in the expression of chronic pain, we tested whether the NAcSh has a similar functional architecture of pain processing stability as we observed in the BLA. We found that light-touch in a SNI state reactivated a greater percentage of NAcSh^*noci*^ ensemble neurons compared to uninjured mice receiving a second *noci*ceptive stimulus. This suggests that the NAcSh^*noci*^ ensemble mirrors the functional stability of the BLA^*noci*^ ensemble and may show an increased recruitment in chronic pain (**Fig. 4K, L**).

**Figure 4.**
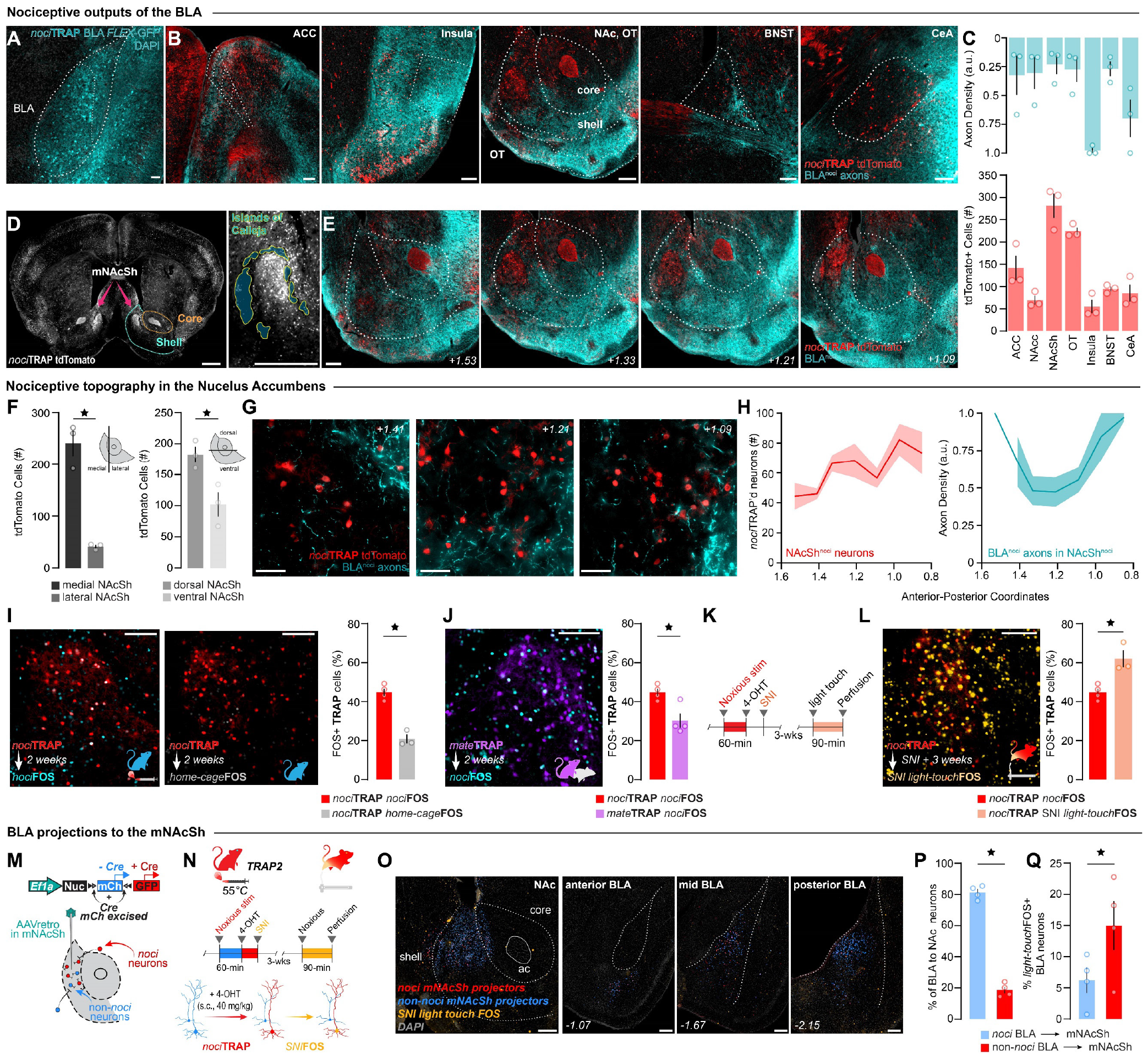
The BLA^*noci*^ ensemble projects to a *noci*ceptive hotspot in the dorsomedial NAcSh. (A) Viral expression of AAV5-hSyn-FLEX-GFP in the right BLA^*noci*^ ensemble of TRAP2 mice. Scale: 100 μm. (B) BLA^*noci*^ ensemble axons (blue) in areas with *noci*ceptive neurons expressing tdTomato (red). Scale: 200 μm. (C) Quantification of BLA^*noci*^ axon density by area (blue) and tdTomato (red) in ipsilateral regions of interest. (D) *noci*TRAP captures a *noci*ceptive hotspot in the dorsomedial NAcSh (NAcSh^*noci*^) encompassed by the Islands of Calleja. Scale: 500 μm. (E) BLA^*noci*^ ensemble axons and NAcSh^*noci*^ neurons spread throughout the anterior-posterior axis of the NAc. Scale: 200 μm. (F) Quantification of tdTomato NAcSh neurons in the medial vs lateral shell (Two-tailed paired t test, p = 0.0115) and in the dorsal vs ventral shell (Two-tailed paired t test, p = 0.0409). (G) 60X images showing BLA^*noci*^ ensemble axons in close proximity to NAcSh^*noci*^ neurons. Scale: 50 μm. (H) Quantification of NAcSh^*noci*^ neurons and BLA^*noci*^ ensemble axon density by area across the anterior-posterior axis of the ipsilateral NAc (n = 3 males). (I) Representative images of *noci*TRAP colocalization with *noci*FOS (blue) vs home-cageFOS (grey) (Two-tailed unpaired t test, p =0.0005, n = 4 *noci*FOS, n = 3 home-cageFOS). Scale: 100 μm. (J) Colocalization of mateTRAP (purple) and *noci*FOS (blue). Scale: 100 μm. Quantification of colocalization of *noci*FOS with neurons captured by mateTRAP relative to *noci*TRAP. (Two-tailed unpaired t test, p =0.0190, n = 4 *noci*TRAP, n = 3 mateTRAP). (K) Timeline of spared nerve injury (SNI) model of chronic neuropathic pain. (L) Colocalization of uninjured *noci*TRAP (red) and light-touch after SNI FOS (orange). Scale: 100 μm. Quantification of colocalization of light-touchFOS in animals than underwent SNI relative to a second acute *noci*ceptive stimulus in uninjured mice. (Two-tailed unpaired t test, p =0.0096, n = 4 *noci*FOS, n = 3 SNI light-touchFOS). (M) Schematic of retrograde tracing in the mNAcSh. (N) Timeline and schematic of retrograde labeling and reactivation of afferents to the mNAcSh. (O) Representative images of the injection site in the mNAcSh and the anterior, mid and posterior BLA. Scale: 200 μm. (P) Quantification of *noci*ceptive and non-*noci*ceptive BLA neurons that project to the mNAcSh. (Two-tailed paired t test, p = 0.0007). (Q) Quantification of light-touchFOS colocalization in *noci*ceptive and non-*noci*ceptive BLA neurons that project to the mNAcSh. (Two-tailed paired t test, p =0.0191, n = 4 (1 male)). BLA = basolateral amygdala, ACC = anterior cingulate cortex, NAc = nucleus accumbens, OT = olfactory tubercle, BNST = bed nucleus of the stria terminalis, CeA = central amygdala, mNAcSh = medial nucleus accumbens shell.

While the BLA^*noci*^ ensemble axons appeared relatively sparse near the NAcSh^*noci*^ neurons, we confirmed that the BLA^*noci*^ ensemble projects to this region by transfecting the medial NAcSh (mNAcSh) of TRAP2 mice with a retrograde color-switch AAV. This technique allowed us to label afferents to the mNAcSh that are *noci*ceptive (*noci*TRAP) with GFP and non-*noci*ceptive afferents with mCherry (**Fig. 4M**). We further labeled chronic pain-active neurons by stimulating the injured hindpaw of SNI mice with a light-touch to evoke FOS (**Fig. 4N-Q, Fig. S5A-C**). Of all BLA neurons, 18.34% project to the mNAcSh; of these projectors, 16.58% are *noci*ceptive, confirming that the BLA→mNAcSh *noci*ceptive projection is a subpopulation of all BLA output to the mNAcSh (**Fig. 4P, Fig. S5D-I**). When we looked at the FOS+ neurons, we found that a significantly larger number of BLA^*noci*^→mNAcSh neurons were reactivated in chronic pain compared to that of the non-*noci*ceptive projection neurons (**Fig. 4Q, Fig. S5J**). This suggests that the stability of the BLA^*noci*^ ensemble across pain states is conserved in the projections to the mNAcSh, indicating this projection is utilized for both acute and chronic pain-related information transfer. Furthermore, while we observed that the BLA^*noci*^ ensemble encompasses the entirety of the anterior-posterior axis of the BLA, the BLA^*noci*^→mNAcSh ensemble is primarily localized to the posterior half of the BLA (**Fig. S5E, F**). Based on the projection target, there is a further subdivision of BLA^*noci*^ neurons, which suggests differential roles of pain processing in anterior *vs*. posterior BLA^*noci*^ neurons that can be refined based on downstream projection target.

### The BLA^noci^ ensemble transmits nociceptive information to the nucleus accumbens

Ca^2+^ recordings from genetically-defined BLA somas retrolabeled from the NAc show increased activity in response to negative valence information, including to noxious foot shock (10). However, given the branching architecture of axonal collaterals and the observation that electric shocks do not fully activate the BLA^*noci*^ ensemble (12), it remains unknown whether the BLA axons transmit information about acute nociceptive stimuli to the NAc, nor how the presence of chronic pain changes this transmission in response to noxious and innocuous stimuli. In fact, no direct BLA axon terminal Ca^2+^ recordings within the NAc have been demonstrated. To synthesize our functional, transcriptomic, and projection-defined characterization of nociceptive processing within the BLA, we recorded BLA axon terminal Ca^2+^ activity in the mNAcSh in response to stimuli across valence and sensory modalities. As our transcriptomic data suggested that the *Rspo*2+ BLA neurons display increased IEG activity in response to noxious stimuli, we utilized *Rspo2*-Cre mice (8) and implanted a fiber optic above the region of the NAcSh*noci*, where we identified a *noci*ceptive hotspot of the NAc (**Fig. 5A, Fig. S6**). Surprisingly, this putative negative valence ensemble displayed decreased Ca^2+^ activity in response to innocuous, aversive, and appetitive stimuli (**Fig. 5B-D**). Our sequencing of the BLA identified two unique subclusters of BLA neurons that express *Rspo2*, containing individual nuclei that show heterogenous responses to noxious stimuli (**Fig. 2M, 3F**). Our axon Ca^2+^ activity recording along with this transcriptomic observation suggests that focusing on a single genetic marker may miss critical information about valence function within the BLA. Therefore, we recorded the Ca^2+^ activity of BLA^*noci*^ ensemble axon terminals in the mNAcSh using the TRAP2 mouse (**Fig. 5E**). The BLA^*noci*^ ensemble axons showed increased Ca^2+^ responses to noxious stimuli across sensory modalities and decreased Ca^2+^ responses to innocuous and appetitive stimuli (**Fig. 5F-H**). This supports previous findings that valence ensembles in the BLA display decreased activity to opposing valence stimuli (8). Finally, we tested the impact of chronic pain on *noci*ceptive transmission from the BLA^*noci*^ ensemble to the mNAcSh (**Fig. 5I**). Interestingly, in a chronic neuropathic pain state, the BLA^*noci*^ ensemble axons show increased Ca^2+^ activity in response to a light touch, suggesting a neural signature of allodynia in the BLA to mNAcSh circuit (**Fig. 5J-L, L, Fig. S7**). We conclude that there are functional subpopulations of *Rspo*2+ BLA neurons important for transmitting *noci*ceptive information to the mNAcSh in acute and chronic pain. Furthermore, examining neural populations based only on single genetic markers, single projection targets, or not considering neural ensembles at distinct anatomical locations within the BLA misses important information for identifying functional subpopulations related to valence processing.

**Figure 5.**
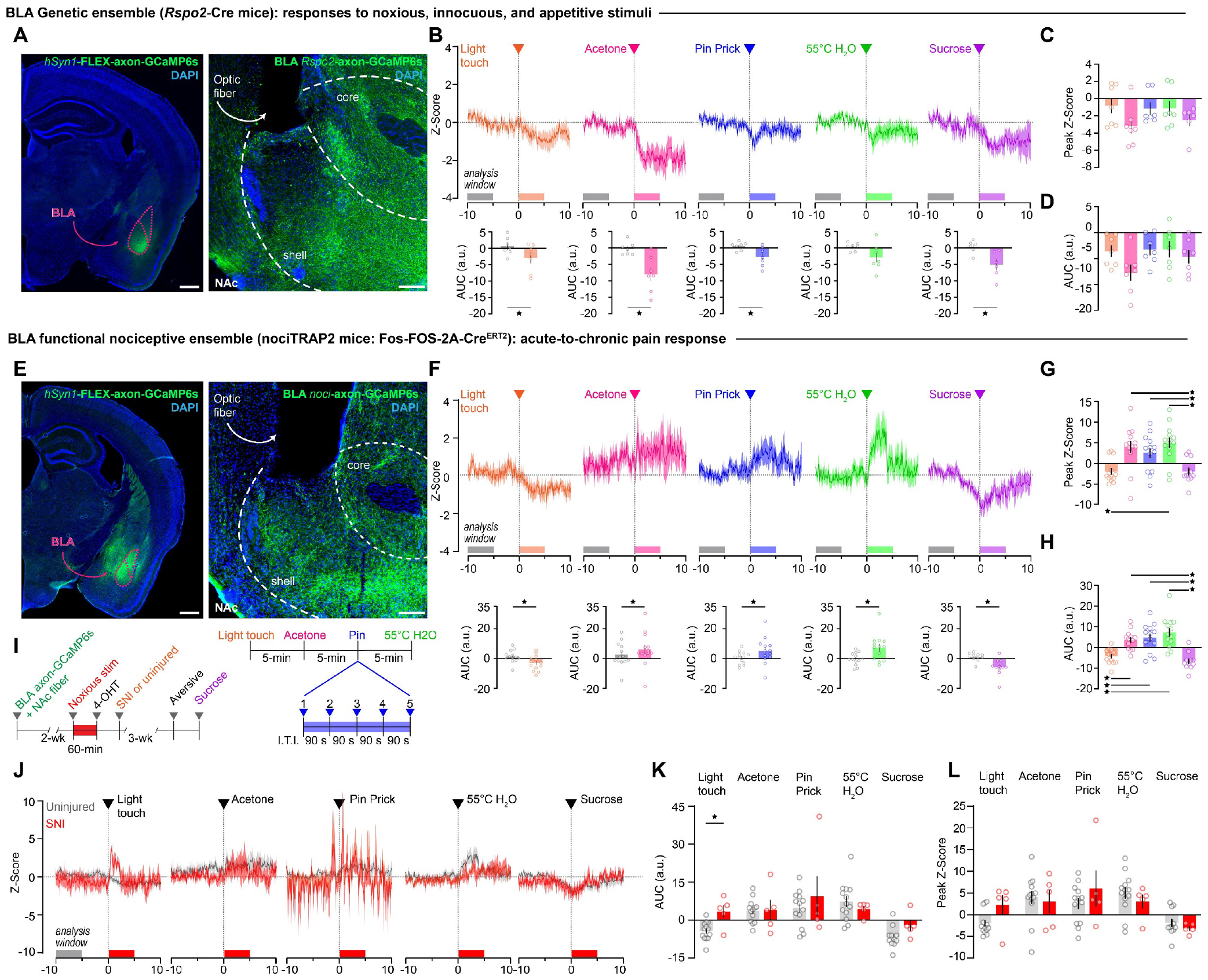
The BLA transmits *noci*ceptive information to the dorsomedial NAcSh. (**A**) Expression of AAV5-hSyn1-FLEX-axon-GCaMP6s in the BLA of *Rspo2*-Cre mice; Expression of axon-GCaMP6s in the NAcSh with an optic fiber placed above the mNAcSh. Scale: 500 μm (left image), 100 μm (right image). (B) BLA *Rspo2* axon terminals in the NAcSh display significantly decreased Ca^2+^ activity in response to innocuous, noxious, and appetitive stimuli compared to pre-stimulus baselines. (Two-tailed paired t test, light touch: p = 0.0196; acetone: p = 0.0027; pin prick: p = 0.0242; 55°C water: p = 0.0849; sucrose: p = 0.0062). (C) Quantification of peak Z-Score of all stimuli. (Mixed-effects analysis, p = 0.2213, n = 7). (D) Quantification of area under the curve (A.U.C) of all stimuli. (Mixed-effects analysis, p = 0.1178; n = 7 (4 males); n = 6 for sucrose. (E) Expression of AAV5-hSyn1-FLEX-axon-GCaMP6s in the BLA*noci* ensemble of TRAP2 mice; Expression of axon-GCaMP6s in the NAcSh with an optic fiber placed above the mNAcSh. Scale: 500 μm (left image), 100 μm (right image). (F) BLA*noci* ensemble axon terminals in the NAcSh display significantly increased Ca^2+^ activity in response to noxious stimuli compared to pre-stimulus baselines. (Two-tailed paired t test, acetone: p = 0.0173; pin prick: p = 0.0288; 55°C water: p = 0.0044, n = 13). In contrast, BLA*noci* ensemble axon terminals display decreased Ca^2+^ activity in response to innocuous and appetitive stimuli compared to pre-stimuli baselines. (Two-tailed paired t test, light touch: p = 0.0013; sucrose: p = 0.0004, n = 13). (G) The peak Z-Score of Ca^2+^ responses to noxious stimuli is significantly different from the response to innocuous and appetitive stimuli. (Mixed-effects analysis with Bonferroni, p = 0.0001; light touch vs. 55°C water: p = 0.004, sucrose vs. acetone: p = 0.0328, sucrose vs. pin prick: p = 0.0566, sucrose vs. 55°C water: p = 0.0133; n = 13 (6 males)). (H) Quantification of A.U.C. of all stimuli. (Mixed-effects analysis with Bonferroni, p < 0.0001; light touch vs. acetone: p = 0.0009, light touch vs. pin prick: p = 0.001, light touch vs. 55°C water: p = 0.0014, sucrose vs. acetone: p = 0.0003, sucrose vs. pin prick: p = 0.0006, sucrose vs. 55°C water: p = 0.0013; n = 13 (6 males)). (I) Timeline for recording Ca^2+^ responses after before and after SNI; stimulation and recording paradigm. (J) BLA*noci* ensemble axons in the NAcSh display increased Ca^2+^ activity in response to a light touch compared to uninjured mice. Ca^2+^ responses to other noxious or appetitive stimuli were unchanged in SNI. (K) The A.U.C. was elevated in response to a light touch in mice with SNI compared to uninjured mice. (Mixed-effects analysis with Bonferroni; main effect of stimulus: p = 0.0009; light touch SNI vs uninjured: p = 0.0388; n = 13 uninjured (6 males), n = 5 SNI (2 males)). (L) The peak Z-Score of the Ca^2+^ responses to stimuli was not significantly different between uninjured and SNI mice. (Mixed-effects analysis with Bonferroni; main effect of stimulus: p = 0.0002; n = 13 uninjured, n = 5 SNI). A.U.C. = area under the curve (arbitrary units) calculated as A.U.C. post-stimulus (0 to 5 seconds) minus A.U.C. pre-stimulus (−10 to -5 seconds). Peak Z-score is calculated over 10 seconds post-stimulus. BLA = basolateral amygdala, NAc = nucleus accumbens.

## Discussion

### The BLA^noci^ ensemble is essential for and stable across pain states

Consistent with our prior single cell Ca^2+^ imaging data and our use of the TRAP1 mouse (12, 85), we report that the BLA contains a stable ensemble of *noci*ceptive neurons (BLA^*noci*^ ensemble) that are non-overlapping with neutral or appetitive valence ensembles. The existence of non-overlapping valence ensembles in the BLA, demonstrated in the current study and by others, raises questions about how the BLA directs attention and affective responses when stimuli of opposing valences are simultaneously present. Future work should examine the contribution of BLA valence ensembles to behavior in opposing motivation behavior tests. In the current study, we extend our prior work to show that acute and chronic pain aversion requires a common BLA^*noci*^ neural ensemble. This suggests that the BLA may be a prime target for non-opioid analgesic drugs to treat acute and chronic pain. Furthermore, inhibition of the BLA^*noci*^ ensemble did not drive appetitive behavior, suggesting treatments that target the BLA^*noci*^ ensemble may have decreased abuse liability compared to opioid analgesics. Therefore, it is critical to examine molecular targets of the BLA^*noci*^ ensemble that can be used for drug targeting.

### Single nuclei RNA sequencing genetically characterizes nociception in the amygdala

To home in on the unique characteristics of the BLA^*noci*^ ensemble, we used snRNAseq to examine the impact of acute and chronic pain on the BLA for the first time. The location of our tissue punches allowed us to capture multiple amygdalar nuclei, including the CeA and the BLA, both of which have been implicated in the processing of pain and valence. Although our current examination focused on the BLA, future analysis on other amygdalar nuclei implicated in pain and negative affect, including the CeA and the ITC will be conducted (56, 58, 86). We found two BLA subclusters (BLA0 and BLA4) that express the gene *Rspo2*, a purported marker of negative valence BLA neurons. While the BLA^*noci*^ ensemble neurons are primarily *Rspo*2+, the majority of *Rspo*2+ neurons are non-*noci*ceptive. Recently, the function of the *Rspo*2+ BLA neurons as a homogenous negative valence population has been challenged. We examined differences in the expression of a panel of 25 IEGs in BLA0 and BLA4 that may account for these observations. Interestingly, nuclei within the BLA0 and BLA4 subclusters displayed diverse IEG activity to a noxious stimulus, suggesting that there are *noci*ceptive subpopulations within the *Rspo*2+ BLA neurons and that *Rspo*2+ BLA neurons are not functionally homogenous. The existence of a functional subpopulation of *Rspo*2+ BLA neurons may rectify the discrepancy between recently published papers on the function of the *Rspo*2+ BLA neurons. Future work will examine the valence and modality specificity of individual *Rspo*2+ BLA neurons. Importantly, our study is the first to examine the impact of *noci*ception on the BLA, identifying multiple genes of interest that should be examined as targets for analgesics.

### Chronic neuropathic pain imparts a transcriptional signature on the BLA

Though discovery of novel analgesics for acute pain would help millions of people, there is a critical need for analgesics that treat debilitating chronic pain. Acute and chronic pain utilize a common BLA^*noci*^ ensemble for pain aversion. Therefore, examination of the effect of chronic pain on the BLA may elucidate mechanisms that can be targeted for acute and chronic pain treatment. Our transcriptional analysis pooling neurons from uninjured and SNI mice identified eight putative BLA subclusters, reflective of the eight putative BLA subclusters when analyzing uninjured mice only. We identified many DEGs between nuclei from uninjured and SNI mice looking at the total BLA and six DEGs when focusing on the *Rspo*2+ BLA0 and BLA4 subclusters. A comparison of the IEG+ nuclei vs IEG-nuclei in BLA0 and BLA4 in SNI conditions revealed only seven DEGs that are altered in a chronic neuropathic state, one of which (*Alkal2*) has been implicated in chronic pain (69). Future investigations will focus on the function of these genes in the BLA in affective pain behavior. Interestingly, when we used a gene ontology analysis of the DEGs unique to a SNI state across all BLA clusters, we found many that are altered by chronic pain and that related to axon regeneration and synaptic transmission. This suggests that chronic pain may change *noci*ceptive transmissions from the BLA to downstream brain regions.

### The BLA^noci^ ensemble projects to the nociceptive hotspot of the NAcSh

Our snRNAseq data suggests that the major molecular changes in the BLA after chronic pain may be on the BLA axons that project to other brain regions. Therefore, we examined the downstream targets of the BLA^*noci*^ axons in brain areas that displayed high *noci*ception-related activity. We observed BLA^*noci*^ axons surrounding a *noci*ceptive hotspot of the NAcSh (NAcSh^*noci*^). This suggests that the NAcSh^*noci*^ neurons may be a downstream target of the BLA^*noci*^ ensemble. The NAcSh^*noci*^ is a previously undescribed functional ensemble of the mNAcSh that is not active at baseline and is not responsive to appetitive stimuli. These neurons are primarily in the posterior medial shell, a region that has been previously described as important for aversion (36, 40, 41, 46). We further characterized this functional ensemble using fluorescent in situ hybridization and determined that the NAcSh^*noci*^ neurons are a mix of dopamine 1 (*Drd1*+) and dopamine 2 receptor (*Drd*2+) neurons (**Fig. S5M**). Interestingly, others have reported the *Drd*2+ neurons of the NAcSh become hyperactive in a chronic pain state (32, 33). This warrants further investigation into the contribution of the *Drd1*+ vs *Drd*2+ NAcSh^*noci*^ neurons to pain-related behaviors. The mNAcSh also receives *noci*ceptive and non-*noci*ceptive projections from many non-amygdalar brain regions, including the ventral tegmental area (VTA) and paraventricular thalamus (PVT) (**Fig. S5K, L**). Future work will examine the contributions of the afferents to the NAcSh^*noci*^, particularly those that densely innervate this region. Interestingly, the majority of BLA^*noci*^ axons in the NAcSh were not in the region where we observed the NAcSh^*noci*^ neurons. This raises the possibility that the BLA^*noci*^ ensemble synapses onto non-*noci*ceptive NAcSh neurons, therefore disinhibiting the NAcSh^*noci*^ neurons. Additional studies are needed to understand the microcircuitry and functional connectivity of the BLA^*noci*^ ensemble axon terminals and NAcSh^*noci*^ neurons.

### The Rspo2+ BLA neurons do not label a single valence projection to the NAcSh

While the BLA projections to the NAc have been primarily studied for their role in positive valence, recent reports suggest that some BLA neurons that project to the NAc encode negative valence information (10, 23). In the current study, we recorded the Ca^2+^ activity of *Rspo*2+ BLA axon terminals in the mNAcSh, where we identified the NAcSh^*noci*^ neurons. Surprisingly, we observed a decrease in axon Ca2+ activity in response to innocuous, noxious, and appetitive stimuli. In concert with our snRNAseq data, this suggests that the *Rspo*2+ BLA neurons are not a homogenous negative valence ensemble. Recent reports found that, in the posterior BLA, *Rspo2* is expressed in putative appetitive neural ensembles (10). Our tracing data suggests that the majority of mNAcSh-projecting BLA neurons are in the posterior BLA. Together, this data indicates that *Rspo*2+ BLA neurons projecting to the mNAcSh may be comprised of a mixed valence population. Therefore, recording Ca2+ activity based on a single genetic marker may not elucidate the function of BLA neurons to valenced stimuli.

### The BLA^noci^ ensemble transmits information to the NAcSh about acute and chronic pain

To coalesce our behavioral, transcriptomic, and anatomical characterization of the *noci*ceptive BLA to NAcSh circuit, we recorded Ca^2+^ activity from BLA^*noci*^ ensemble axons in the mNAcSh in response to innocuous, noxious, and appetitive stimuli. Interestingly, we observed decreased axon Ca^2+^ activity in response to innocuous and appetitive stimuli. This may suggest that other BLA ensembles that project to the mNAcSh are engaged in response to non-noxious salient stimuli and that *noci*ceptive information transmission is suppressed in safe conditions. Previous reports suggest that activity of BLA valence ensembles is antagonistic—increased activity of one is correlated with decreased activity of the other (8, 11). Further work is needed to elucidate the mechanism of inhibition of BLA valence ensembles. The BLA^*noci*^ ensemble axons in the mNAcSh displayed increased Ca2+ activity in response to noxious stimuli across sensory modalities. Increased Ca2+ activity within BLA^*noci*^ axon terminals suggests that there is downstream activation of the mNAcSh in response to noxious stimuli. Our work is the first to show *noci*ceptive information transmission from the BLA to the NAcSh and adds to the growing challenges against the dogma that this circuit primarily functions to transmit appetitive-related valence signals.

### The BLA^noci^ ensemble transmits a signature of allodynia to the NAcSh

Our prior work imaging single cell Ca2+ activity in the BLA revealed that the BLA^*noci*^ ensemble is stable across the development of chronic pain. Furthermore, we showed that the BLA^*noci*^ ensemble expands its response profile in chronic pain, showing increased Ca2+ activity to light touch, which it does not respond to in an uninjured state. This activity correlates with increased pain behavior to light touch after SNI (allodynia). In the current study, we extended these findings, showing that there is increased BLA^*noci*^ ensemble axon terminal Ca2+ activity in response to a light touch following SNI in the NAcSh, activity that mirrors the allodynia behavior we observe in mice with SNI. This coupled with the increased activation of NAcSh^*noci*^ neurons following SNI suggests that this neural circuit from the BLA^*noci*^ ensemble to the NAcSh may be particularly important in chronic pain. Further work will examine the impact of manipulating this circuit, including the downstream NAcSh^*noci*^ neurons, in acute and chronic pain.

In all, for the first time, we have described a *noci*ceptive projection from the BLA to the mNAcSh that is valence-specific, active to noxious stimuli regardless of sensory modality, and stable across acute and chronic pain sates. Moreover, the BLA to NAcSh neural circuit may play a distinct role in the allodynia that is a signature of chronic neuropathic pain. Our results further support the BLA as a prime target for novel analgesics to treat acute and chronic pain and emphasize the importance of examining the downstream connectivity of the BLA with other affective brain regions.

Acknowledgments.

We thank the University Laboratory Animal Resources (ULAR) group at the University of Pennsylvania for assistance with rodent husbandry and veterinary support at the Translational Research Laboratory building. We thank Clinton Wojick for assistance with data processing and analysis. We would also like to thank other members of the Corder Lab, Maxx Yung, Adrienne Jo, Gregory Salimando, and Justin James for additional technical assistance and support. We thank Susumu Tonegawa for providing the *Rspo2*-Cre mice.

## Funding

National Institutes of Health NIGMS DP2GM140923 (GC)

National Institutes of Health NIDA R00DA043609 (GC)

National Institutes of Health NINDS F31NS125927 (JAW)

National Institutes of Health NIMH DP2MH129985 (EK)

Linda Pechenik Montague Investigator Award (EK)

National Institutes of Health NIEHS T32ES019851 (AP)

National Institutes of Health NIDA F32DA055458 (BAK)

National Institutes of Health NIDA F32DA053099 (NMM)

National Institutes of Health NIDA F31DA057795 (LMW)

National Institutes of Health NIDA R21DA057458 (RCC)

National Institutes of Health NIDA R21DA055846 (BCR)

## Author contributions

*Conceptualization*: JAW, GC

*Immunohistochemistry, in situ hybridization*: JAW, MM, JKC, CSO, LMW, NMM, LLE

*Mouse intracranial surgeries*: JAW, CSO

*Mouse behavior, analyses, and statistics*: JAW, GC, BK, JKC, AH, NMM, LMW, LLE

*RNA sequencing –tissue collection*: JAW, BK

*RNA sequencing – nuclei isolation*: SNC, BCR

*RNA sequencing - data analysis*: AP, RCC, BCR, EK

*Writing original draft*: JAW, GC

*Review and editing of manuscript*: all authors

## Materials and Methods

### Animals

All experimental procedures were approved by the Institutional Animal Care and Use Committee of the University of Pennsylvania and performed in accordance with the US National Institutes of Health (NIH) guidelines. Male and female mice aged 2-5 months were housed 2-5 per cage and maintained on a 12-hour reverse light-dark cycle in a temperature and humidity-controlled environment. All experiments were performed during the dark cycle. Mice had ad-libitum food and water access throughout experiments. For behavioral, Ca2+ imaging, anatomical, and transcriptomic experiments, we utilized Fos-FOS-2A-iCre^ERT2^ or “TRAP2” mice (Jackson Laboratory, stock #030323) (47) bred to homozygosity. Additional anatomical experiments utilized TRAP2 mice crossed with Ai9 (B6.Cg-Gt(ROSA)26Sortm9(CAG-tdTomato)Hze/J) reporter mice that express a tdTomato fluorophore in a Cre-dependent manner (“TRAP2:tdTomato”) purchased from Jackson Laboratory, stock #007909 bred to homozygosity for both genes. Additional Ca2+ imaging experiments utilized *Rspo2*-Cre mice from Dr. Susumu Tonegawa’s lab (8).

### Viral Vectors

All viral vectors were aliquoted and stored at -80°C until use and then stored at 4°C for a maximum of four days. For optogenetic inhibition experiments, we intracranially injected 400 nL of AAV1-*hSyn1*-SIO-stGACR2-FusionRed (Addgene 105677-AAV1; titer: 1.0 x 1012 vg/mL) or AAV1-*CAG*-FLEX-tdTomato (Addgene 28306-AAV1; titer: 1.0 x 1012 vg/mL) into bilateral BLA at coordinates anteriorposterior (AP): -1.20 mm, mediolateral (ML): ± 3.20 mm, dorsoventral (DV): -5.20 mm. For anatomical tracing experiments, we intracranially injected 400 nL of AAV5-*hSyn*-DIO-EGFP (Addgene 50457-AAV5; titer: 1.3 x 1012 vg/mL) into the right BLA at coordinates AP: -1.20 mm, ML: 3.20 mm, DV: -5.20 mm. For Ca2+ imaging in fiber photometry experiments, we intracranially injected 400 nL of AAV5-*hSyn*-FLEx-axon-GCaMP6s (Addgene 112010-AAV5, titer: 2.2 x 1012 vg/mL) into the right BLA at coordinates AP: -1.20 mm, ML: 3.20 mm, DV: -5.20 mm. For retrograde tracing of BLA inputs to the ACC, we intracranially injected 350 nL of AAVrg-*hSyn*-mCherry (Addgene 114472-AAVrg; titer: 2.4 x 1012 vg/mL) at coordinates AP: 1.50 mm, ML: 0.30 mm, DV: -1.50 mm. For retrograde tracing of BLA inputs to the NAcSh, we intracranially injected 350 nL of AAVrg-*hSyn*-EGFP (Addgene 50465-AAVrg; titer: 2.4 x 1012 vg/mL) into the right NAcSh at coordinates AP: 1.00 mm, ML: 0.60 mm, DV: -4.40 mm. For retrograde tracing of inputs to the NAcSh, we intracranially injected 300 nL of AAV5-*EF1a*-Nuc-flox(mCherry)-EGFP (Addgene 112677-AAV5; titer:) into the right NAcSh at coordinates: AP: 1.20, ML: 0.60, DV: -4.55). For chemogenetic inhibition experiments(87), we intracranially injected 300 nL of AAV9-*hSyn*-DIO-hM4Di(Gi)- mCherry or AAV9-*hSyn*-DIO-mCherry (Addgene 50459-AAV9; titer: 2.33 x 1012 vg/mL) into bilateral NAcSh at coordinates AP: 1.2, ML: ± 0.6 mm, DV: -4.55 mm.

### Stereotaxic surgery

Adult mice (∼8 weeks of age) were anesthetized with isoflurane gas in oxygen (initial dose = 5%, maintenance dose = 1.5%), and fitted into WPI or Kopf stereotaxic frames for all surgical procedures. 10 µL Nanofil Hamilton syringes (WPI) with 33 G beveled needles were used to intracranially infuse AAVs into the BLA, NAcSh, or ACC. The following coordinates were used, based on the Paxinos mouse brain atlas, to target these regions of interest: BLA (from Bregma, AP: -1.20 mm, ML: ± 3.20 mm, DV:

−5.20 mm), NAcSh (from Bregma, AP: 1.20 mm, ML: ± 0.60 mm, DV:−4.55 mm), ACC (from Bregma, AP: -1.50 mm, ML: ± 0.3 mm, DV:−1.5 mm). Mice were given a 3–8-week recovery period to allow ample time for viral diffusion and transduction to occur. For fiber photometry studies, following viral injection into the right BLA, we placed a fiberoptic implant (5.2 mm fiber, 400 µm diameter, Doric Lenses) approximately 0.2 mm above the DV coordinate of the injection site for the NAcSh. After setting the fiberoptic in position, MetaBond (Parkell) and Jet Set dental acrylic (Lang Dental) were applied to the skull of a mouse to rapidly and firmly fix the fiberoptic in place. In brief, after exposure of the skull, the bone was scored with a scalpel blade and a small skull screw (∼1.7 mm diameter, 1.6 mm length) was placed to provide grooves for the MetaBond to provide additional adhesion points. The MetaBond reagent was liberally applied over the skull and up and along the fiberoptic cannula. Once dried, the MetaBond was then covered with a layer of Jet Set acrylamide to create a reinforced head cap, as well as to cover the exposed skin of the incision site. Mice were then given a minimum of 3 weeks to recover and allow for optimal viral spread and transduction along axon terminals prior to beginning in vivo Ca2+ recordings. For all surgical procedures in mice, meloxicam (5 mg/kg) was administered subcutaneously at the start of the surgery, and a single 0.25 mL injection of sterile saline was provided upon completion. All mice were monitored for up to three days following surgical procedures to ensure the animals’ proper recovery and to provide additional daily subcutaneous meloxicam.

### Chronic neuropathic pain model

As described previously (12), to induce a chronic pain state, we used a modified version of the Spared Nerve Injury (SNI) model of neuropathic pain, as previously described (70). This model entails surgical section of two of the sciatic nerve branches (common peroneal and tibial branches) while sparing the third (sural branch). Following SNI, the receptive field of the lateral aspect of the hindpaw skin (innervated by the sural nerve) displays hypersensitivity to tactile and cool stimuli, eliciting pathological reflexive and affective-motivational behaviors (allodynia). To perform this peripheral nerve injury procedure, anesthesia was induced and maintained throughout surgery with isoflurane (4% induction, 1.5% maintenance in oxygen). The left hind leg was shaved and wiped clean with alcohol and betadine. We made a 1-cm incision in the skin of the mid-dorsal thigh, approximately where the sciatic nerve trifurcates. The biceps femoris and semimembranosus muscles were gently separated from one another with blunt scissors, thereby creating a <1-cm opening between the muscle groups to expose the common peroneal, tibial, and sural branches of the sciatic nerve. Next, ∼2 mm of both the common peroneal and tibial nerves were transected and removed, without suturing and with care not to distend the sural nerve. The leg muscles are left uncultured and the skin was closed with tissue adhesive (3M Vetbond), followed by a Betadine application. During recovery from surgery, mice were placed under a heat lamp until awake and achieved normal balanced movement. Mice were then returned to their home cage and closely monitored over the following three days for well-being.

### TRAP protocol (tamoxifen induction)

#### NociTRAP

*Noci*TRAP induction was performed similarly to previously described (12). We habituated mice to a testing room for two to three consecutive days. During these habituation days, no *noci*ceptive stimuli were delivered and no baseline thresholds were measured (i.e. mice were naïve to pain experience before the TRAP procedure). We placed individual mice within red plastic cylinders (∼9-cm D), with a red lid, on a raised perforated, flat metal platform (61-cm x 26-cm). The experimenter’s sat in the testing room for the thirty minutes of habituation; this was done to mitigate potential alterations to the animal’s stress and endogenous anti*noci*ception levels. To execute the TRAP procedure, we placed mice in their habituated cylinder for 30 min, and then a 55°C water droplet was applied to the central-lateral plantar pad of the left hindpaw) once every 30 s over 10 min. Following the water stimulations, the mice remained in the cylinder for an additional 60 min before injection of 4-hydroxytamoxifen (4-OHT, 40 mg/kg in vehicle; subcutaneous). After the injection, the mice remained in the cylinder for an additional 4 hours to match the temporal profile for c-FOS expression, at which time the mice were returned to the home cage.

#### Home-cageTRAP

Home-cageTRAP induction was performed without habituation. At least 2 hours into the dark cycle, mice were gently removed from their home cages. Mice were then injected with 4-OHT (40 mg/kg in vehicle; subcutaneous) and returned to their home cages.

#### MateTRAP

The process of mateTRAP began with habituation of mice to “home away from home” (HAFH) cages for one hour a day for four consecutive days beginning at least 2 hours into the dark cycle. During these habituation sessions, animals were isolated in HAFH cage and no stimuli were delivered. On the fifth day, mice were placed alone in HAFH cages for one hour, then were administered 4-OHT (40 mg/kg in vehicle; subcutaneous) before being gently placed back in HAFH cages. Immediately after injection, an age-matched, mating-receptive (estrus or proestrus phase of estrus cycle) conspecific of the opposite sex was introduced to the HAFH cage for four hours to match the temporal profile for 4-OHT. During the social interaction, a video recording was taken to allow for post-hoc manual quantification of mounting behaviors. After that, mice were returned to their home cages.

### Immunohistochemistry

Animals were anesthetized using FatalPlus (Vortech Pharamaceuticals) and transcardially perfused with 0.1 M phosphate buffered saline (PBS), followed by 10% normal buffered formalin solution (NBF, Sigma, HT501128). Brains were quickly removed and post-fixed in 10% NBF for 24 hours at 4 °C, and then cryo-protected in a 30% sucrose solution made in 0.1 M PBS until sinking to the bottom of their storage tube (∼48 h). Brains were then frozen in Tissue Tek O.C.T. compound (Thermo Scientific), coronally sectioned on a cryostat (CM3050S, Leica Biosystems) at 30 μm or 50 μm and the sections stored in 0.1 M PBS. Floating sections were permeabilized in a solution of 0.1 M PBS containing 0.3% Triton X-100 (PBS-T) for 30 min at room temperature and then blocked in a solution of 0.3% PBS-T and 5% normal donkey serum (NDS) for 2 hours before being incubated with primary antibodies (1°Abs included: chicken anti-GFP [1:1000, Abcam, ab13970], guinea pig anti-FOS [1:1000, Synaptic Systems, 226308], rabbit anti-FOS [1:1000, Synaptic System, 226008],rabbit anti-DsRed [1:1000, Takara Bio, 632496]; prepared in a 0.3% PBS-T, 5% NDS solution for ∼16 h at room temperature. Following washing three times for 10 min in PBS-T, secondary antibodies (2°Abs included: Alexa-Fluor 647 donkey anti-rabbit [1:500, Thermo Scientific, A31573], Alexa-Fluor 488 donkey anti-chicken [1:500, Jackson Immuno, 703-545-155], Alexa-Fluor 555 donkey anti-rabbit [1:500, Thermo Scientific, A31572] Alexa-Fluor 647 donkey anti-guinea pig [1:500, Jackson Immuno, 706-605-148], prepared in a 0.3% PBS-T, 5% NDS solution were applied for ∼2h at room temperature, after which the sections were washed again three times for 5 mins in PBS-T, then again three times for 10 min in PBS-T, and then counterstained in a solution of 0.1 M PBS containing DAPI (1:10,000, Sigma, D9542). Fully stained sections were mounted onto Superfrost Plus microscope slides (Fisher Scientific) and allowed to dry and adhere to the slides before being mounted with Fluoromount-G Mounting Medium (Invitrogen, 00-4958-02) and cover slipped.

### Fluorescence in situ hybridization

Animals were anesthetized using isoflurane gas in oxygen, and the brains were quickly removed and fresh frozen in O.C.T. using Super Friendly Freeze-It Spray (Thermo Fisher Scientific). Brains were stored at −80° C until cut on a cryostat to produce 16 μm coronal sections of the NAcSh. Sections were adhered to Superfrost Plus microscope slides, and immediately refrozen before being stored at −80° C. Following the manufacturer’s protocol for fresh frozen tissue for the V2 RNAscope manual assay (Advanced Cell Diagnostics), slides were fixed for 15 min in ice-cold 10% NBF and then dehydrated in a sequence of ethanol serial dilutions (50%, 70%, and 100%). Slides were briefly air-dried, and then a hydrophobic barrier was drawn around the tissue sections using a Pap Pen (Vector Labs). Slides were then incubated with hydrogen peroxide solution for 10 min, washed in distilled water, and then treated with the Protease IV solution for 30 min at room temperature in a humidified chamber. Following protease treatment, C1 and C2 cDNA probe mixtures specific for mouse tissue were prepared at a dilution of 50:1, respectively, using the following probes from Advanced Cell Diagnostics: *Fos* (C1, 316921), *Drd2* (C2, 406501-C2), and *Drd1* (C3, 406491-C3). Sections were incubated with cDNA probes (2 h), and then underwent a series of signal amplification steps using FL v2 Amp 1 (30 min), FL v2 Amp 2 (30 min) and FL v2 Amp 3 (15 min). 2 min of washing in 1x RNAscope wash buffer was performed between each step, and all incubation steps with probes and amplification reagents were performed using a HybEZ oven (ACD Bio) at 40° C. Sections then underwent fluorophore staining via treatment with a serious of TSA Plus HRP solutions and Opal 520, 570, and 620 fluorescent dyes (1:5000, Akoya Biosystems, FP1487001KT, FP1495001KT). All HRP solutions (C1-C2) were applied for 15 min and Opal dyes for 30 min at 40° C, with an additional HRP blocker solution added between each iteration of this process (15 min at 40° C) and rinsing of sections between all steps with the wash buffer. Lastly, sections were stained for DAPI using the reagent provided by the Fluorescent Multiplex Kit. Following DAPI staining, sections were mounted, and cover slipped using Fluoromount-G mounting medium and left to dry overnight in a dark, cool place. Sections from all mice were collected in pairs, using one section for incubation with the cDNA probes and another for incubation with a probe for bacterial mRNA (dapB, ACD Bio, 310043) to serve as a negative control.

### Imaging and Quantification

All tissue was imaged on a Keyence BZ-X all-in-one fluorescent microscope at 48-bit resolution using the following objectives: PlanApo-λ x4, PlanApo-λ x20 and PlanApo-λ x40. All image processing prior to quantification was performed with the Keyence BZ-X analyzer software (version 1.4.0.1). Quantification of neurons expressing fluorophores was performed via manual counting of TIFF images in Photoshop (Adobe, 2021) using the Counter function or using HALO software (Indica Labs), which is a validated tool for automatic quantification of fluorescently-labeled neurons in brain tissue (88–90). Counts were made using 20X magnified z-stack images of a designated regions of interest (ROI). For axon density quantification, immunohistochemistry was performed to amplify the GFP signal and visualize ipsilateral BLA^*noci*^ axons throughout the brain in 50 μm tissue free floating slices as described above. Areas with dense axon innervation were identified using 4X imaging. Areas implicated in emotion and *noci*ception were selected for additional 20X imaging with Z-stacks. These regions of interest (ROIs) were initially visualized at 20X to determine the region with the highest fluorescence. The exposures for FITC and CY3 were adjusted to avoid overexposed pixels for the brightest area. This exposure was kept consistent for all slices for an individual mouse. For an individual ROI, one slice per mouse was included. ROIs were drawn in ImageJ and threshold intensity was measured. The maximum intensity was set to a value of 1.0 and all ROIs are reported as intensity compared to the maximum.

### Single nuclei RNA sequencing

#### Nuclei preparation

A single punch of the right side of the BLA measuring 2 mm in width and 1 mm in depth, was used to prepare the nuclei suspensions. Nuclei isolation was performed utilizing the Minute™ single nucleus isolation kit designed for neuronal tissue/cells (Cat# BN-020, Invent Biotechnologies). Briefly, the tissue was homogenized using a pestle in a 1.5 mL LoBind Eppendorf tube. Subsequently, the cells were resuspended in 700 µl of cold lysis buffer and RNAse inhibitor and incubated on ice for 5 minutes. The homogenate was then transferred to a filter within a collection tube and incubated at -20°C for 8 minutes. Following this, the tubes were centrifuged at 13,000 x g for 30 seconds, the filter was discarded, and the samples were centrifuged at 600 x g for 5 minutes. The resulting pellet underwent one wash with 200 µL of PBS + 5% BSA and then resuspended in 60 µL of PBS + 1% BSA. The concentration of nuclei in the final suspension was assessed by staining with Trypan Blue and counting using a hemacytometer. The suspension was diluted to an optimal concentration of 500-1000 nuclei/µL.

#### Single-Nuclei Gene Expression Assay

The single-nuclei gene expression assay (snRNAseq) was conducted following the instructions provided by 10x Genomics. A total of 20,000 nuclei were loaded into the 10x Genomics microfluidics Chromium controller, with the aim of recovering approximately 10,000-12,000 nuclei per sample. Subsequently, sequencing libraries were constructed following the manufacturer’s protocol for the Chromium Next GEM Single Cell 3ʹ v3.1 kit. Libraries containing individual and unique indexes were pooled together at equimolar concentrations of 1.75 nM and sequenced on the Illumina NovaSeq 6000, using 28 cycles for Read 1, 10 cycles for the i7 index, 10 cycles for the i5 index, and 90 cycles for Read 2.

### Data analysis

#### RNA sequencing

##### Preprocessing of scRNAseq data

Paired end sequencing reads were processed using 10x Genomics Cellranger v5.0.1. Reads were aligned to the mm10 genome optimized for single cell sequencing through a hybrid intronic read recovery approach (91). In short, reads with valid barcodes were trimmed by TSO sequence, and aligned using STAR v2.7.1 with MAPQ adjustment. Intronic reads were removed and high-confidence mapped reads were filtered for multimapping and UMI correction. Empty GEMs were also removed as part of the pipeline. Initial dimensionality reduction and clustering was performed prior to processing to enable batch correction and removal of cell free mRNA using SoupX (92).Raw expression matrices with counted, individual nuclei UMI and genes were used for subsequent steps and filtering by QC metrics.

#### Clustering and merging by condition and comparison

Raw matrices for each individual replicate per condition were converted to Seurat objects using Seurat 5.0.1, and filtered to remove UMIs with thresholds of > 200 minimum features, < 5% mitochondrial reads, and < 5% ribosomal reads. Replicates were merged to generate objects per condition for the subsequent steps. Each dataset was normalized using the default scale factor of 10000, variable selection was performed using 2000 features, then scaled using default parameters. Dimensionality reduction with PCA used the first 30 principal components and the nearest-neighbor graph construction used the first 10 dimensions. Clustering was next performed using a resolution of 0.4 before layers corresponding to each replicate were integrated using CCAIntegration with a k weight of 60 and then rejoined. The dataset per condition was then dimensionally reduced using the integrated CCA at with 30 dimensions and the same resolution of 0.4. ScType (93) was used for automated, de novo cell type identification of the clusters followed by manual curation for clusters with low confidence scores. For all comparisons, the objects per condition were merged and processed using the same integration methodology above to scale and normalize between all incorporated samples. Non-neuronal cell types were excluded from all downstream analysis steps after initial classification. Nuclei expressing the top 75th percentile of Slc17a7 (Vglut1) expression were marked as the basolateral amygdala (BLA). The inhibitory cell clusters were marked by the top 75th percentile of either Gad1 or Gad2 expression, and no detected Slc17a7 or Slc17a6 (Vglut2) transcripts. Each independent dataset was then subclustered with a resolution of 0.4. BLA and inhibitory cluster separated objects were then merged, normalized, and dimensionally reduced to generate the BLA-Inhibitory clusters dataset. All differentially expressed genes were identified using FindMarkers with the Wilcoxon Rank Sum test with Bonferroni correction for multiple testing with a minimum pct of 0.25 and fold change threshold of 1.2. Genes with an adjusted p-value of < 0.01 were considered differentially expressed.

#### Modular activity scoring and subsetting

Modular activity scores were calculated for glutamatergic neuron and BLA datasets using AddModuleScore with the list of the 25 putative immediate early genes (Arc, Bdnf, Cdkn1a, Dnajb5, Egr1, Egr2, Egr4, Fos, Fosb, Fosl2, Homer1, Junb, Nefm, Npas4, Nr4a1, Nr4a2, Nr4a3, Nrn1, Ntrk2, Rheb, Sgsm1, Syt4, Vgf) against a control feature score of 5 (67). Nuclei in the top 90th activity score percentile were isolated as IEG+ while those under the 50th percentile threshold were IEG-. The merged object was aggregated and dimensionally reduced with a resolution of 0.2. FindMarkers was used to perform all IEG+/IEG-pairwise comparisons.

#### Gene ontology analysis

Gene ontology (GO) analyses were performed for DEGs between uninjured and SNI conditions for all BLA subclusters. GO analysis was performed via PANTHER (protein analysis through evolutionary relationships) v 18.0 on genes that were upregulated in the SNI conditions using the Statistical enrichment test with Bonferroni correction and a p-value cut off of 0.05. The list of DEGs was assessed for biological processes, molecular function, and cellular components.

### Behavioral tests

On test days, mice were brought into procedure rooms ∼1 h before the start of any experiment to allow for acclimatization to the environment. Mice were provided food and water ad libitum during this period. For multi-day testing conducted in the same procedure rooms, animals were transferred into individual “home away from home” (HAFH) cages ∼1 h prior to the start of testing and were only returned to their home cages at the end of the test day. All testing and acclimatization were conducted under red light conditions (<10 lux). Equipment used during testing was cleaned with a 70% ethanol solution before starting, and in between, each behavioral trial to mask odors and other scents.

### Sensory testing for pain affective-motivational and nociceptive reflex behavioral assays

To evaluate mechanical reflexive sensitivity, we used a logarithmically increasing set of 8 von Frey filaments (Stoelting), ranging in gram force from 0.07-to 6.0-g. These filaments were applied perpendicular to the plantar hindpaw with sufficient force to cause a slight bending of the filament. A positive response was characterized as a rapid withdrawal of the paw away from the stimulus within 4 seconds. Using the Up-Down statistical method, the 50% withdrawal mechanical threshold scores were calculated for each mouse and then averaged across the experimental groups (12).

To evaluate affective-motivational responses evoked by mechanical stimulation, we used a sharp 25G syringe needle (pin prick) (12). The pin prick was applied as a sub-second poke to the hindpaw, and the duration of attending behavior was collected for up to 30 s after the stimulation.

To evaluate affective-motivational responses evoked by thermal stimulation (12), we applied either a single, unilateral 55°C drop of water or acetone (evaporative cooling) to the left hindpaw, and the duration of attending behavior was collected for up to 30 s after the stimulation. Only one drop stimulation was applied on a given testing session. Additionally, we used an inescapable hotplate set to 50° C. The computer-controlled hotplate (6.5 in x 6.5 in floor, Bioseb) was surrounded by a 15 in high clear plastic chamber and two web cameras were positioned at the front or side of the chamber to continuously record animals to use for post hoc behavioral analysis. For the tests conducted for chemogenetic activation studies, mice were administered CNO 30 min prior to the start of behavior testing to allow for complete absorption of the drug and previous sensory testing fd(87). Mice were gently placed in the center of the hotplate floor and removed after 60 seconds.

### Conditioned place avoidance

Fifteen minute pre- and post-conditioning test sessions and three twice daily 30-min conditioning sessions consisting of alternating side-pairings of CNO (3.0 mg/kg, i.p.) and saline (0.9% saline, i.p.) were used to determine conditioned place preference (Walters et al., 2005). During the pre-test session, mice were placed in a two-chambered place preference chamber (Med Associates, inc., St. Albans, VT) inside a sound-attenuated chamber (Med Associates, inc., St. Albans, VT) and allowed to explore both sides for 15 min (900 s). The amount of time spent on each side was recorded, and data were used to assign the animals to be conditioned in the non-preferred chamber. Mice were conditioned twice daily for 3 days following the pre-test, with the saline-paired box conditioning sessions in the AM and the CNO-paired conditioning sessions in the PM, separated by 4 hours. Each group received CNO (3 mg/kg) on one side and saline (0.9% sodium chloride) on the other. Locomotor activities for each conditioning session were measured using two consecutive beam breaks. One day after the last conditioning session, animals were allowed to explore freely between the two sides for 15 min (900s), and time spent on each side was recorded. The preference score (time spent on the drug-paired side minus time in the saline-paired side on the post-conditioning day minus the preconditioning day) was calculated for each mouse.

### In vivo fiber photometry calcium recordings and data analysis

Optical recordings of axon-GCaMP6s fluorescence were acquired using an RZ10x fiber photometry detection system (Tucker-Davis Technologies), consisting of a processor with Synapse software (Tucker-Davis Technologies), and optical components (Doric Lenses and ThorLabs). Excitation wavelengths generated by LEDs (460 nm blue light and 405 nm violet light) were relayed through a filtered fluorescence minicube at spectral bandwidths of 460–495 and 405 nm to a pre-bleached, low auto-fluorescence, mono fiberoptic patch cord connected to the implant on top of each animal’s head. Power output for the primary 460 nm channel at the tip of the fiberoptic cable was adjusted to ∼100 µW. Signal in both 460 and 405 nm channels was monitored continuously throughout all recordings, with the 405 nm signal used as an isosbestic control for both ambient fluorescence and motion artifacts introduced by movement of components in the light path. Wavelengths were modulated at frequencies of 210– 220 and 330 Hz, respectively. All signals were acquired at 1 kHz and lowpass filtered at 4 Hz. On testing days, mice were connected to the photometry system, and following a 10 min habituation period, recording sessions began. Each stimulus type was applied 5 times, with an inter-trial interval (ITI) of 90 seconds. Between different stimulus types (0.07 g VF filament, acetone droplet, 25G needle pick prick, 55°C water droplet), the LED was turned off for 5 minutes to minimize photobleaching. Following testing, all mice were perfused, and the tissue was assessed for proper viral targeting and transduction efficacy, as well as optic fiber placement via immunohistochemistry (see above section for details).

Analysis of the GCaMP signal was performed with the use of the open source, fiber photometry analysis MATLAB software suite, pMAT (94). Using pMAT, bulk fluorescent signal from both the 460 and 405 channels were normalized to compare differences in calcium-mediated event metrics for both the total duration of a recording (frequency) and at select events using peri-event time histogram (PETH) analyses locked to specific behaviors designated by the application of an external transistor-transistor logic (TTL) input. Amplitude and area under the curve were determined from PETH analyses across groups. The MATLAB polyfit function was used to correct for the bleaching of signal for the duration of each recording, using the slope of the 405 nm signal fitted against the 460 nm signal. Detection of GCaMP-mediated fluorescence is presented as a change in the 460 nm/fitted 405 nm signal over the fitted 405 signal (ΔF/F). From ΔF/F values, robust Z-scores were calculated for analysis (94).

### In vivo optogenetic inhibition

At the time of bilateral virus injection, mice were implanted with bilateral optic fibers (200 μm diameter, Prizmatix) above the BLA. The mice were left to recover for 2 weeks before *noci*TRAP and 3 weeks after *noci*TRAP to ensure ample virus expression. Optogenetic stimulation was delivered with a blue LED (455 nm) via a bifurcated fiber optic cable. Power output at the tip was measured to be ∼ 5 mW per side. For sensory testing, mice were placed on the elevated wire-floor von Frey rack in small plexiglass containers for 30 minutes. Sensory testing was conducted before LED stimulation, during a 30-second stimulation, and after stimulation with a 90 second ITI.

### Drugs

4-hydroxytamoxifen (4-OHT; Hellobio, catalog #HB6040) was dissolved in 100% ethanol on the morning of use. The solution was further diluted in Kolliphor EL (Sigma Aldrich, catalog #C5135-500G) and finally 1X phosphate buffered saline (PBS; Calbiochem, catalog #524650). Clozapine N-oxide (CNO dihydrochloride, water soluble; HelloBio HB6149) was delivered i.p. at a dose of 3.0 mg/kg body weight.

### Statistics and data presentation

The number of animals used in each experiment were pre-determined based on analyses of similar experiments in the literature and supplemented as needed based on observed effect sizes. When male and female mice were used (all experiments except the RNA-sequencing), an analysis for sex differences in the data was performed when powered to do so. When no significant effect of sex was observed, data was presented with male and female mice combined. In some figures, mice or data points were removed due to experimenter or technological errors. All data are presented as mean ±the standard error of the mean (SEM) for each group, and all statistical analyses were performed using Prism 9 & 10 software (GraphPad Software). Data was represented using Prism and changed aesthetically using Illustrator (Adobe). Representative images of histology were selected to display viral spread, fiber optic placements, and endogenous fluorescence. Where applicable, we only adjusted histology images with linear manipulations of contrast and brightness.

## Supplemental Figures

**Supp. Fig. 1.**
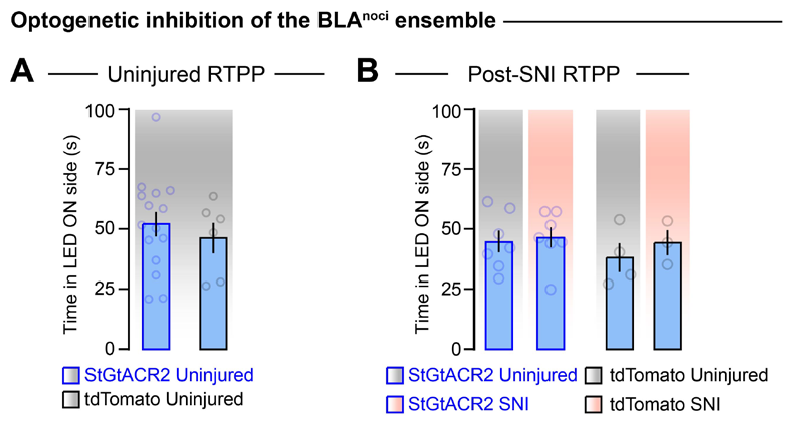
Inhibition of the BLA^noci^ ensemble does not significantly decrease hyperalgesia. (A) Inhibition of the BLA^noci^ ensemble does not induce a place preference in uninjured (Two-tailed unpaired t test, p = 0.5294, n = 15 StGtACR2 (7 male), n = 6 tdTomato (2 male)) or (B) SNI mice (Two-way ANOVA with Bonferroni, interaction: p = 0.6786, n = 7 StGtACR2 uninjured (4 male), n = 7 StGtACR2 SNI (3 male), n = 4 tdTomato uninjured (2 male), n = 3 tdTomato SNI (all female)).

**Supp. Fig. 2.**
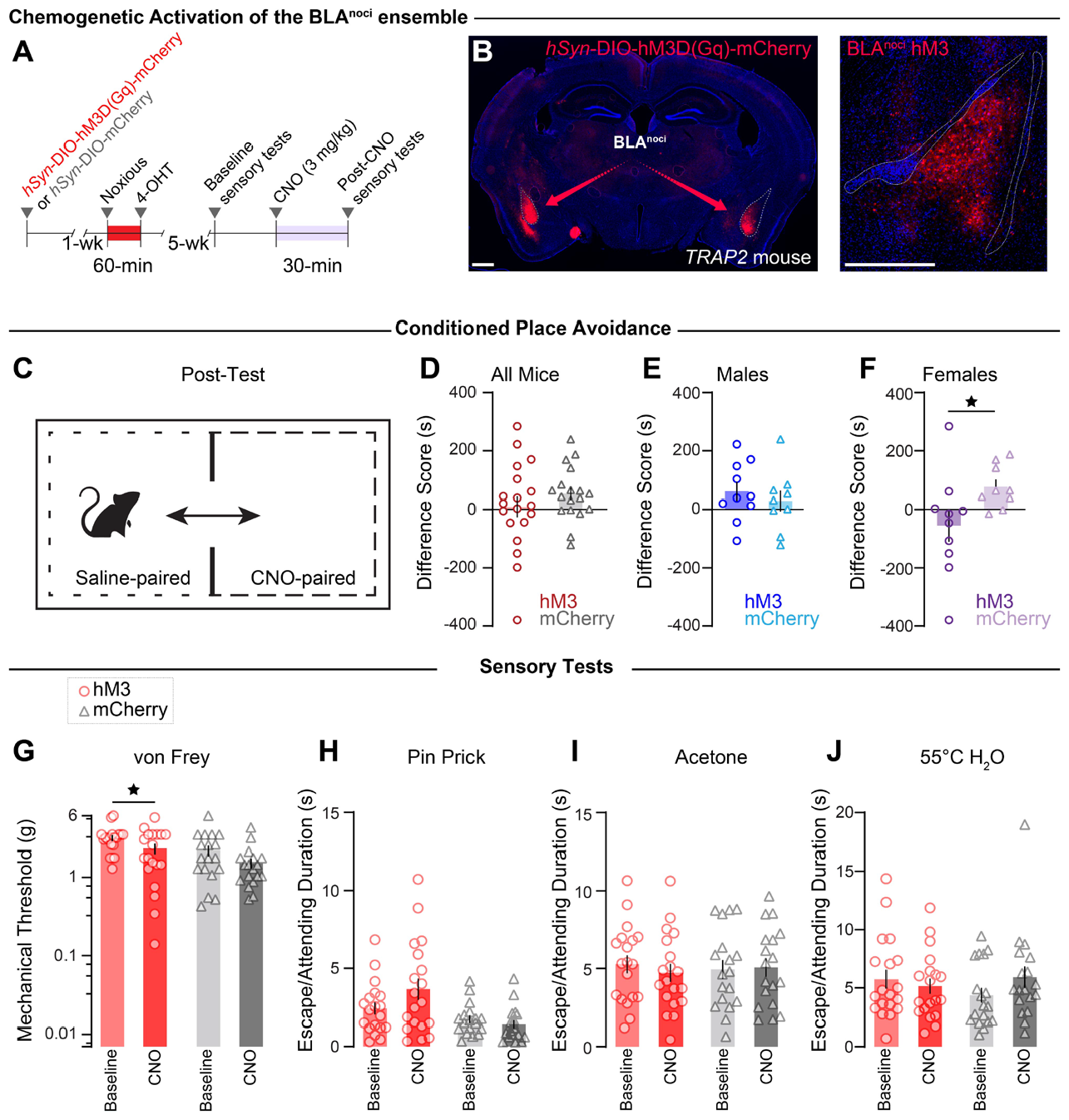
Activation of the BLA^noci^ ensemble is not sufficient to produce hyperalgesia. (A) Timeline for chemogenetic activation of BLA^noci^ ensemble. (B) Expression of AAV5-*hSyn*-DIO-hM3D(Gq)-mCherry bilaterally in BLA^noci^ neurons. Scale: 500 μm. (C) Conditioned place avoidance assay. (D) Male and female mice combined do not display a CPA (Two-tailed unpaired t test, p = 0.2949), (E) nor do males alone (Two-tailed unpaired t test, p = 0.5044), (F) but female mice display a CPA to BLA^noci^ ensemble activation (Two-tailed unpaired t test, p = 0.0496, n = 10 hM3). (G) Chemogenetic activation of the BLA^noci^ ensemble decreased mice’s mechanical threshold in the von Frey assay (Two-way ANOVA with Bonferroni, main effect of drug: p = 0.0049, and virus: p = 0.0148; hM3 baseline vs. CNO: p = 0.0469). (H-J) Chemogenetic activation of the BLA^noci^ ensemble did not impact nocifensive behavioral responses to noxious stimuli (Two-way ANOVA with Bonferroni: pin prick: main effect of virus: p = 0.0021; acetone: interaction: p = 0.5429; 55°C water: interaction: p = 0.1057; n = 19 hM3 (9 male), n = 18 mCherry (9 male)). BLA = basolateral amygdala.

**Supp. Fig. 3.**
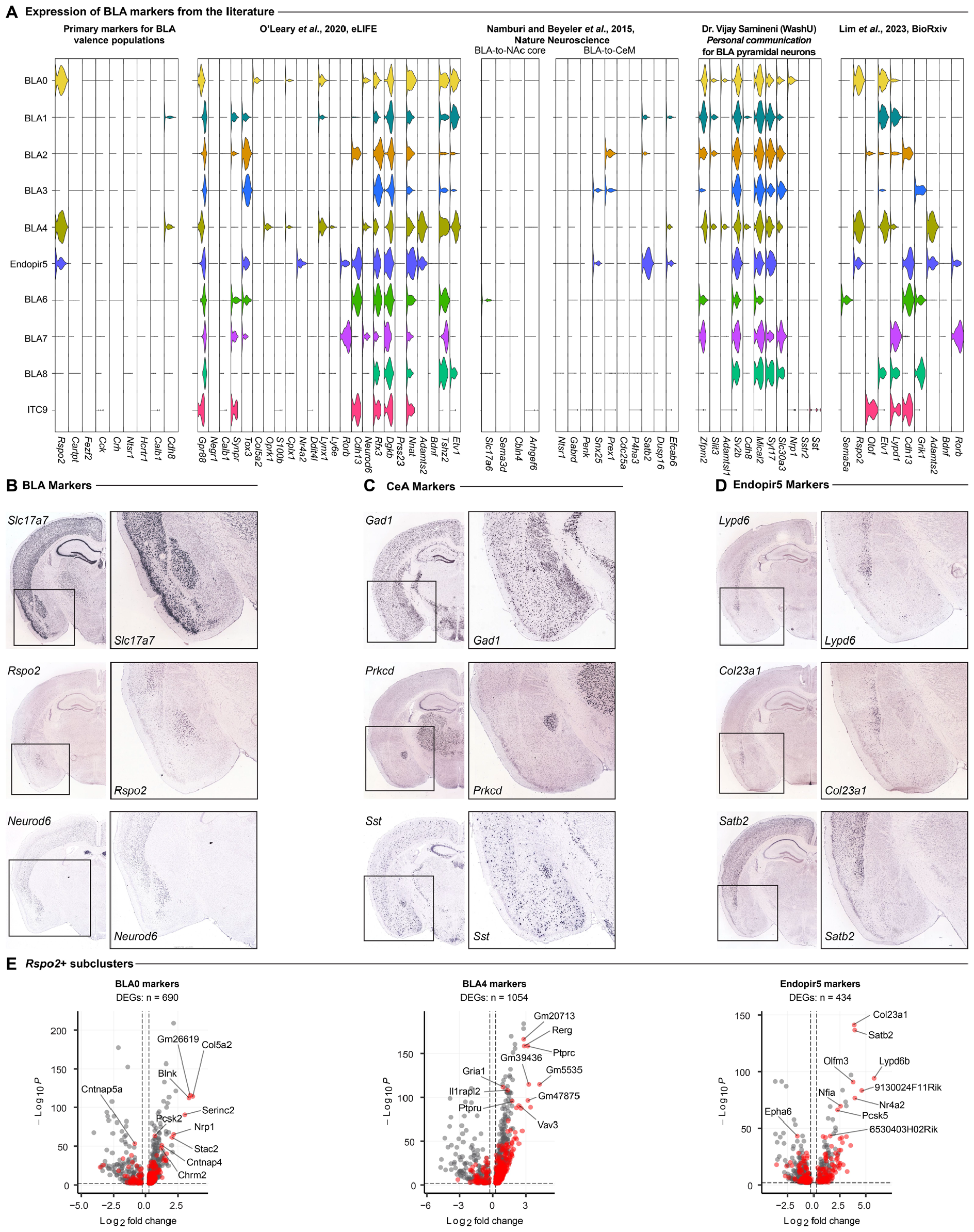
The 10 BLA subclusters express previously published BLA genes at varying levels. (A) Expression levels of genes reported to mark BLA ensembles across various published, pre-printed, and personally discussed works. (B-D) *In situ* hybridization images from the Allen Brain mouse atlas displaying multiple genes enriched in each cluster that defines the BLA, CeA, or the Endopiriform cortex. (E) DEGs unique to each *Rspo2*+ subcluster initially defined as BLA nuclei. Red dots indicate genes enriched in that condition and grey dots shared DEGs across conditions.

**Supp. Fig. 4.**
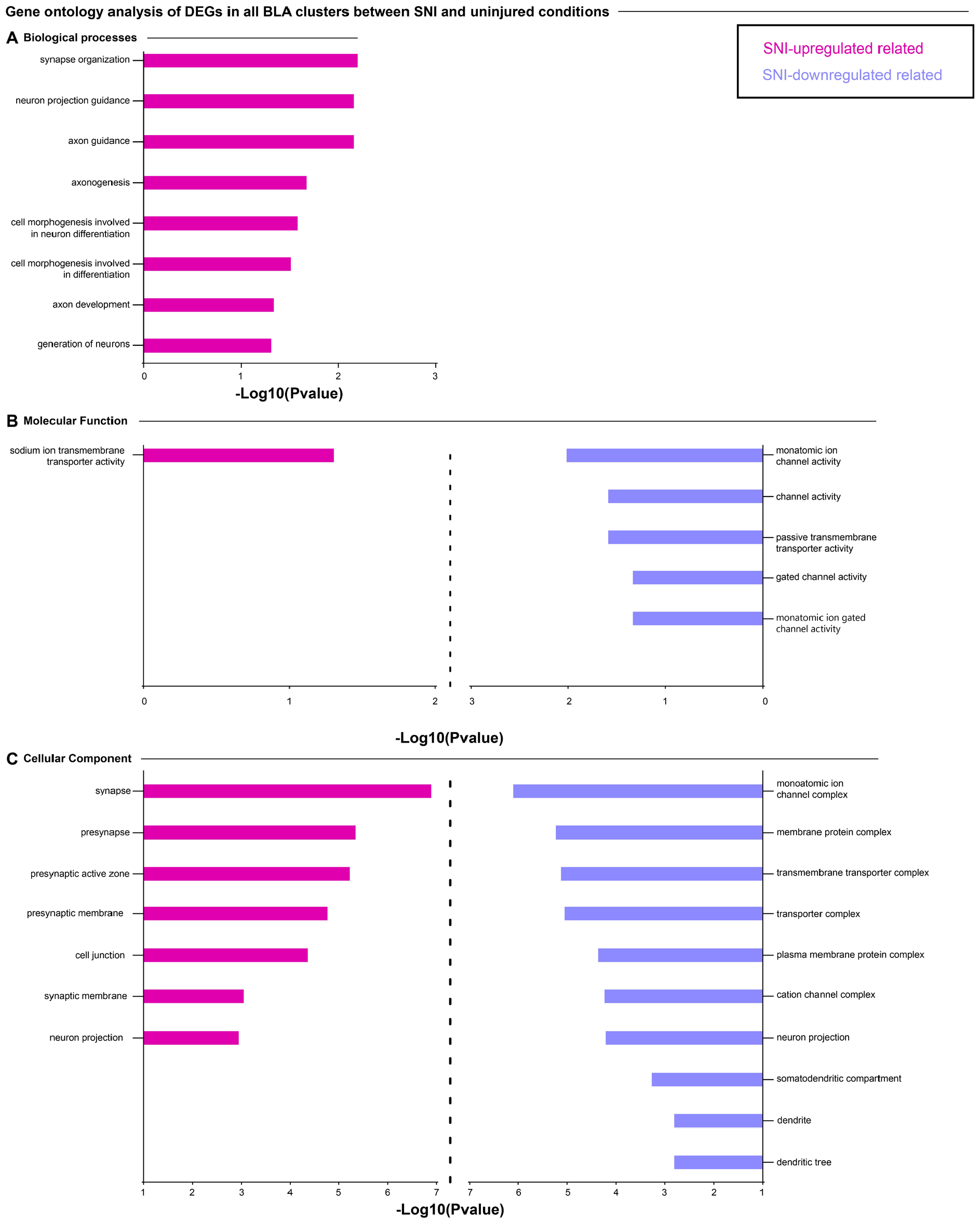
SNI results in transcriptional disruptions to the *Rspo2*+ BLA subclusters. (A) Gene ontology analysis of DEGs from Fig. 4D specific to the effect of SNI on the 8 putative BLA subclusters in regards to biological processes, (B) molecular functions, and (C) cellular components. Pink bars represent analyses of genes that were upregulated in SNI conditions, purple bars represent analyses of genes that were downregulated in SNI conditions.

**Supp. Fig. 5.**
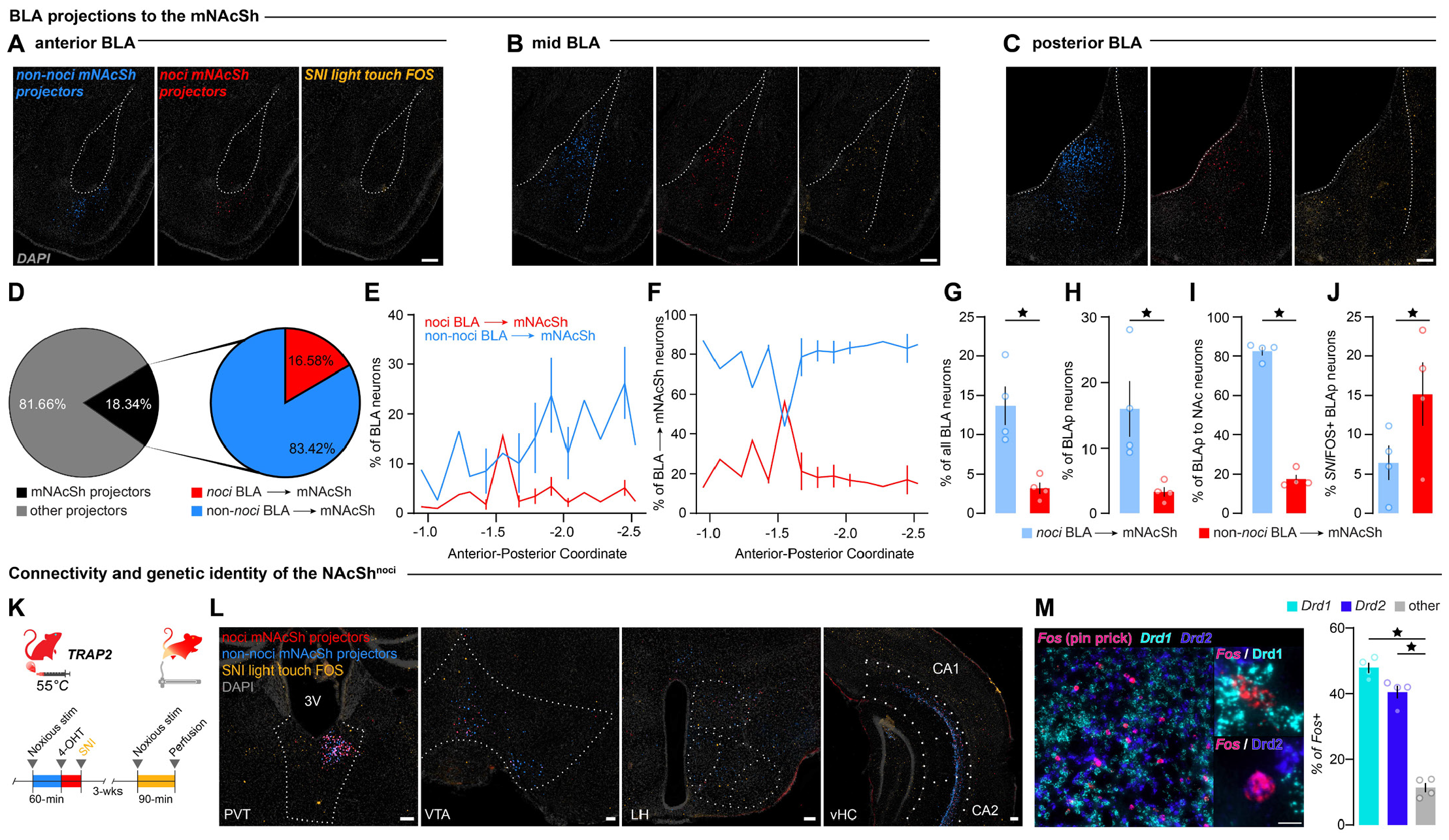
The mNAcSh receives nociceptive and non-nociceptive inputs from affective brain regions. (A-C) Individual channels showing non-nociceptive (blue) and nociceptive (red) BLA neurons that project to the mNAcSh and BLA neurons that express FOS in response to a light touch in SNI (yellow). (D) Quantification of BLA projections to the mNAcSh and BLA→mNAcSh nociceptive vs non-nociceptive projection neurons (E, F) across the anterior-posterior axis. (G) There are significantly more non-nociceptive BLA neurons that project to the mNAcSh across the entire (Two-tailed paired t test, p =0.0112, n = 4) and (H, I) posterior BLA (Two-tailed paired t test, p = 0.0398, 0.0006, n = 4). (J) The nociceptive posterior BLA projections to the mNAcSh are reactivated in a chronic pain state. (Two-tailed paired t test, p =0.0184, n = 4). (K, L) The mNAcSh receives nociceptive and non-nociceptive projections from other affective, nociceptive brain regions. Scale: 100 μm. PVT = paraventricular thalamus, 3V = 3rd ventricle, VTA = ventral tegmental area, LH = lateral hypothalamus, vHC = ventral hippocampus. (M) The mNAcSh^noci^ ensemble is a heterogenous mix of *Drd1*+ and *Drd2*+ and unidentified neurons. (Repeated measured one-way ANOVA, p = 0.001; Drd1 vs other: p = 0.0014; Drd2 vs other: p = 0.0074). Scale: 1 μm. *Drd1* = dopamine 1 receptor mRNA, *Drd2* = dopamine 2 receptor mRNA.

**Supp. Fig. 6.**
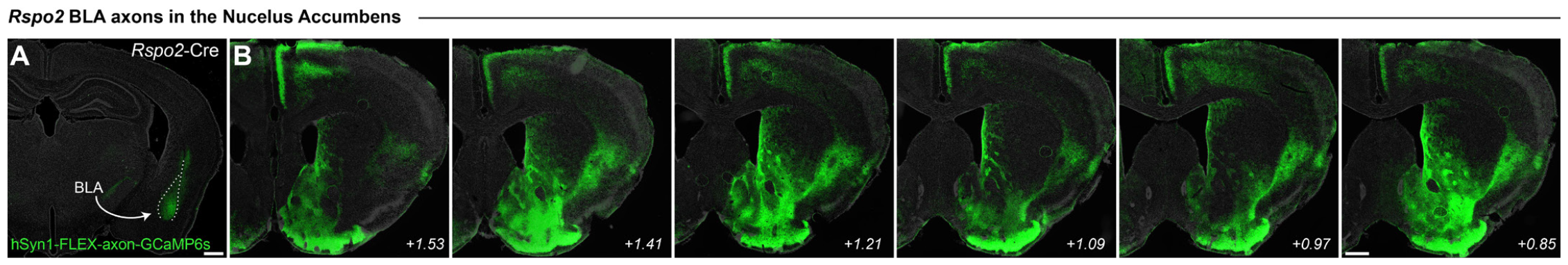
*Rspo2* BLA neurons send dense projections to the NAc. (A) Expression of AAV5-*hSyn1*-FLEX-axon-GCaMP6s in the *Rspo2*+ neurons of the right BLA. Scale: 500 μm. (B) The *Rspo2*+ BLA axons appear throughout the NAc, including around the NAcSh^noci^. Scale: 500 μm. BLA = basolateral amygdala.

**Supp. Fig. 7.**
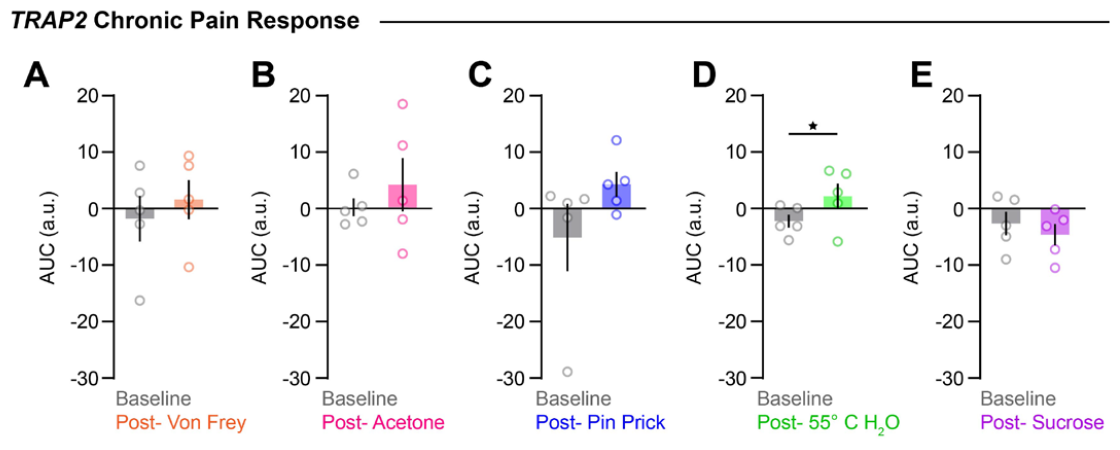
The BLAnoci ensemble responds to noxious stimuli post-SNI. (A-E) Compared to baseline, the BLA^noci^ ensemble shows increased Ca^2+^ activity to innocuous and noxious stimuli post-SNI. (Two-tailed paired t test, Light touch: p = 0.2653, Acetone: p = 0.3675, Pin prick: p = 0.0304, 55°C Water: p = 0.023, Sucrose: p = 0.42, n = 5).

